# Fermented Botanical Product Modulates Soil Bacterial Communities and Enhances Plant-Growth-Promoting Activity for Sustainable Agriculture

**DOI:** 10.64898/2026.05.29.728655

**Authors:** Yukina Adachi-Oshima, Ayano Hojo, Yuri Mizuno, Yusuke Tateuchi, Kotaro Fujioka, Hideto Torii, Yukihiro Tashiro

## Abstract

Although biostimulants have attracted attention for sustainable agricultural systems, their efficacy remains poorly understood. In this study, we evaluated the effects of fermented botanical product (FBP) produced by fermenting and aging 41 types of fruits, grains, seaweed, and root vegetables with brown sugar for more than three years. Three crops, tomato, rice, and komatsuna (*Brassica rapa*), were cultivated with the application of 5,000- or 10,000-fold diluted FBP in greenhouses or fields. Application of diluted FBP promoted plant growth, as indicated by increased fresh weights of shoots, leaves, and roots, fruit production in tomato, and rice husk yield. As diluted FBP contained low nutrient levels, an indirect mechanism of plant growth promotion was suggested. Bacterial community structure analysis indicated changes in alpha diversity, beta diversity, and the predominant phyla in FBP-applied soils without plants and in soils cultivated with tomato, rice, and komatsuna. In addition, the abundance of plant-growth-promoting bacteria, such as *Arthrobacter*, *Pseudomonas*, *Paraburkholderia*, and *Planifilum*, increased in soils treated with diluted FBP. Furthermore, ammonium formation activity was observed in komatsuna cultivation soils treated with diluted FBP, whereas phosphate-solubilizing activity was enhanced in soils from all three crop cultivation systems treated with diluted FBP. These results suggest that diluted FBP influences bacterial communities and promotes crop growth through indirect effects, including increases in plant-growth-promoting bacteria, ammonium production, and phosphate solubilization. Alternatively, FBP may directly stimulate plant growth. Therefore, FBP may be a useful biostimulant for sustainable agricultural systems.

**Highlights:**

- Diluted FBP promoted the growth of tomato, rice, and komatsuna (*Brassica rapa*).
- Diluted FBP altered the bacterial community structure in cultivated soils.
- FBP increased the abundance of plant-growth-promoting bacteria in cultivated soils.
- FBP stimulated ammonium formation and phosphate solubilization in cultivated soils.
- FBP may be a useful biostimulant for sustainable agricultural systems.

**Graphical Abstract:** 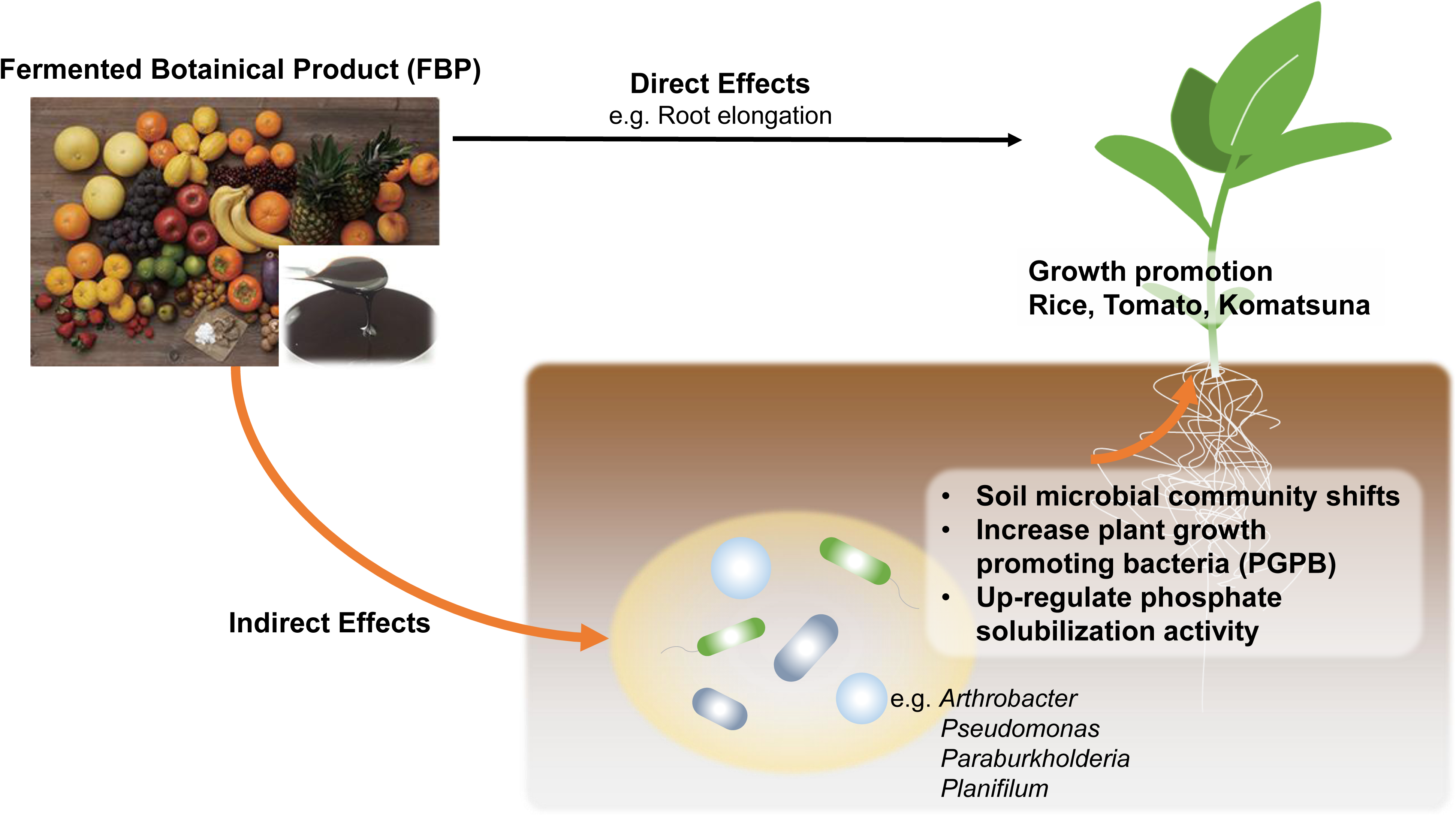

## 1. INTRODUCTION

In the 20th century, agricultural methods using chemical fertilizers primarily composed of nitrogen, phosphorus, and potassium were developed and widely adopted. As fertilizer use increased, agricultural yields rose, contributing to growth of the world population. According to the United Nations report *World Population Prospects*, the global population is projected to reach approximately 9.7 billion by 2050, from approximately 8 billion at present (https://population.un.org/wpp/). Furthermore, among the Sustainable Development Goals (SDGs) established by the United Nations in 2016 (https://sdgs.un.org/partnerships), Goal 2, “End hunger, achieve food security and improved nutrition, and promote sustainable agriculture,” was proposed. Chemical fertilizers are produced using finite resources and large amounts of energy, and their prices are increasing [1]. Their long transport distances have also been shown to contribute substantially to greenhouse gas emissions [2]. Furthermore, dependence on chemical fertilizers causes soil degradation, including erosion, soil compaction, carbon loss, and reduced soil microbial diversity [3,4]. Therefore, establishing and promoting sustainable agricultural methods that do not rely on conventional chemical fertilizers is an urgent task from the perspectives of food supply and environmental conservation.

In Japan, the Ministry of Agriculture, Forestry and Fisheries established the “MIDORI Strategy for Sustainable Food Systems,” which aims to enhance stakeholder engagement at each stage of food supply chains and promote innovation to reduce environmental load, in 2021 (https://www.maff.go.jp/e/policies/env/env_policy/midori.html). The MIDORI Strategy sets targets that include a 30% reduction in chemical fertilizer use and a 25% increase in organic farmland by 2050. Similarly, the European Union (EU) has set a target to reduce chemical fertilizer use by 30% by 2030 [5]. The use of biostimulants is expected to contribute to reductions in chemical fertilizer use. A biostimulant is generally defined as “any substance or microorganism applied to plants with the aim to enhance nutrition efficiency, abiotic stress tolerance and/or crop quality traits, regardless of its nutrient content, excluding pesticides and fertilizers” [6]. Biostimulants include humic and fulvic acids, beneficial fungi and bacteria, and botanicals, among others [6]. Although various biostimulants are being manufactured, the scientific basis for their use and their actual efficacy remain unclear in many cases. Biostimulants contain multiple components, making it difficult to identify the active ingredients. Therefore, clarifying their functional mechanisms is essential for expanding the market for biostimulant products [7].

Fermented botanical product (FBP) is an organic material produced by fermenting and aging 41 plant species for three years. Viable microorganisms are unlikely to remain in the final product. FBP is typically applied after dilution at ratios ranging from 5,000- to 10,000-fold. Previous study has shown that soil drenching with FBP suppresses wilt symptoms in tomato caused by the bacterial wilt pathogen *Ralstonia pseudosolanacearum* [8], suggesting its potential for reducing pesticide use. However, no academic reports have described its plant growth-promoting effects. Therefore, this study aimed to verify the plant growth-promoting effects of FBP in multiple plant species and elucidate its growth-promoting mechanisms. We conducted 1) cultivation tests using tomato, rice, and komatsuna (*Brassica rapa*; Japanese mustard spinach), 2) analyses of bacterial community structure and bacterial counts, and 3) measurements of plant growth-promoting activity.

## 2. MATERIALS AND METHODS

### 2.1. Composition of fermented botanical product

FBP is an organic material produced by Manda Fermentation Co., Ltd. (Japan). FBP is manufactured by fermenting and aging 41 types of fruits, grains, seaweed, and root vegetables with brown sugar for more than three years. During fermentation and aging, no heating, water addition, or microbial seed culture additives are used. A preliminary experiment showed that PCR amplification using a universal eubacterial primer set was not detected in DNA extracted from FBP, suggesting that bacteria were unlikely to remain in FBP. FBP is a dark brown, slightly acidic paste with a pH of approximately 4. Table 1 shows the components of FBP, which particularly contains the following amino acids: valine (Val), lysine (Lys), serine (Ser), proline (Pro), alanine (Ala), leucine (Leu), tyrosine (Tyr), methionine (Met), histidine (His), threonine (Thr), arginine (Arg), glutamate (Glu), isoleucine (Ile), tryptophan (Trp), phenylalanine (Phe), and aspartate (Asp).

**Table 1.** Component of Fermented Botanical Product.

| Component | Unit | Amount per 100 g |
| --- | --- | --- |
| Water | g/100 g | 5.0–50.0 |
| Protein | g/100 g | 0.5–10.0 |
| Lipid | g/100 g | 0.05–10.00 |
| Carbohydrate | g/100 g | 30.0–75.0 |
| Glucose | g/100 g | 20–21 |
| Fructose | g/100 g | 19–20 |
| Sodium | mg/100 g | 20–300 |
| Magnesium | mg/100 g | 40–200 |
| Calcium | mg/100 g | 50–900 |
| Total nitrogen | mg/100 g | 290 |
| Total phosphorus (P<sub>2</sub>O<sub>5</sub>) | mg/100 g | 110 |
| Total potassium (K<sub>2</sub>O) | mg/100 g | 700 |
| Total silica | mg/100 g | <100 |
| Total calcium (CaO) | mg/100 g | 70 |
| Total magnesium (MgO) | mg/100 g | 90 |
| Alkali content | mg/100 g | 200 |
| Total manganese (MnO) | mg/100 g | <1 |
| Total boron (B<sub>2</sub>O<sub>3</sub>) | mg/100 g | <3 |
| Total iron | mg/100 g | 12 |
| Total copper | mg/100 g | 0.4 |
| Total zinc | mg/100 g | 0.4 |
| Total sulfur (SO<sub>3</sub>) | mg/100 g | 380 |
| Total molybdenum | mg/100 g | <1 |
| Total cobalt | mg/100 g | <1 |
| Organic carbon | g/100 g | 27.61 |
| Isoleucine | mg/100 g | 30–200 |
| Leucine | mg/100 g | 50–400 |
| Lysine | mg/100 g | 20–200 |
| Methionine | mg/100 g | 10–100 |
| Phenylalanine | mg/100 g | 30–250 |
| Tyrosine | mg/100 g | 20–200 |
| Threonine | mg/100 g | 40–200 |
| Tryptophan | mg/100 g | 1–100 |
| Valine | mg/100 g | 30–300 |
| Histidine | mg/100 g | 10–200 |
| Arginine | mg/100 g | 40–400 |
| Alanine | mg/100 g | 50–300 |
| Aspartic acid | mg/100 g | 100–600 |
| Glutamic acid | mg/100 g | 100–1200 |
| Glycine | mg/100 g | 30–300 |
| Proline | mg/100 g | 40–400 |
| Serine | mg/100 g | 30–300 |
| Ammonium thiocyanate | mg/100 g | <10 |
| Arsenic | mg/100 g | <0.1 |
| Nitrite (HNO<sub>2</sub>) | mg/100 g | <10 |
| Biuret nitrogen (N) | mg/100 g | <10 |
| Sulfamic acid | mg/100 g | <10 |
| Cadmium | mg/100 g | <0.05 |
| Nickel | mg/100 g | <0.3 |
| Chromium | mg/100 g | <0.3 |
| Titanium | mg/100 g | <5 |
| Mercury | mg/100 g | 0.002 |
| Lead | mg/100 g | <0.3 |

### 2.2. Soil immersion test without plant cultivation

Mixed soil consisting of black soil (Plantation Iwamoto Co., Ltd., Ibaraki, Japan), red soil (Nafco Co., Ltd., Fukuoka, Japan), and humus (E.H Green Corporation, Fukuoka, Japan) at a volume ratio of 2:3:2 was filled into a 72-well tray (n = 4). The mixed soils were immersed for at least 1 h three times per week in purified water (WAT), 200-fold diluted brown sugar solution (SUG), or 100-fold diluted FBP solution (FBP) by bottom watering, basically until the soil surface became moist. When surface moisture was not maintained, water or solution was supplied from the top surface. The trays were incubated at 25 °C for 2 weeks in a glass chamber at the Environmental Control Center for Experimental Biology, Kyushu University.

### 2.3. Tomato cultivation test

Tomato cultivation was conducted at 25 °C for 112 days from May to August 2023 using a completely randomized design with five replicates in a glass chamber at the Environmental Control Center for Experimental Biology, Kyushu University. The treatments consisted of water (WAT) or 5,000-fold diluted FBP (FBP) application, with 0% (zero), 50% (half), or 100% (full) chemical fertilizer. The reagents used were 228 and 456 mg NH₄NO₃ (Nacalai Tesque, Kyoto, Japan), 891.5 and 1783 mg CaH₄P₂O₈ (Katayama Chemical, Osaka, Japan), 324 and 648 mg KNO₃ (Sigma-Aldrich, Tokyo, Japan), and 203 and 406 mg MgSO₄·7H₂O (FUJIFILM Wako Pure Chemical Corporation, Osaka, Japan) for the 50% and 100% NPK fertilizer applications, respectively. In addition, 52.5 and 105 mg NH₄NO₃ (Nacalai Tesque), 205.5 and 411 mg CaH₄P₂O₈ (Katayama Chemical), and 74.5 and 149 mg KNO₃ (Sigma-Aldrich) were applied for the 50% (half) and 100% (full) NPK fertilizer treatments, respectively. These reagents were mixed with unsterilized soil as basal fertilizers 1 week before seed sowing. A mixture of 214 g black soil (Plantation Iwamoto Co., Ltd.), 270 g red soil (Nafco Co., Ltd.), and 70 g humus (E.H Green Corporation) per pot was sieved through an 8 mm stainless steel sieve. Approximately 500 g of mixed soil was placed in each 0.7 L pot (130 mm in diameter and 113 mm in height). During cultivation, 1 mL of 5,000-fold diluted FBP was applied weekly. Agronomic variables, including leaf number, fresh weight of leaves and shoots, and fresh biomass, were recorded at harvest.

### 2.4. Rice cultivation test

Rice cultivation was conducted at 25 °C for 129 days from May to September 2023 using a completely randomized design with five replicates in a glass chamber at the Environmental Control Center for Experimental Biology, Kyushu University. The treatments consisted of water (WAT) or 5,000-fold diluted FBP (FBP) application, with 0% (zero), 50% (half), or 100% (full) chemical fertilizer. The reagents used were 236 and 472 mg NH₄NO₃ (Nacalai Tesque), 400 and 800 mg CaH₄P₂O₈ (Katayama Chemical), 111 and 222 mg KCl (Sigma-Aldrich), and 218 and 436 mg Mg₂Si₃O₈·5H₂O (FUJIFILM Wako Pure Chemical Corporation, Osaka, Japan) for the 50% and 100% NPK fertilizer applications, respectively. In addition, 43 and 86 mg NH₂CONH₂ (FUJIFILM Wako Pure Chemical Corporation) were applied as the 50% (half) and 100% (full) NPK fertilizer treatments, respectively, after 95 days of cultivation. These reagents were mixed with unsterilized soil as basal fertilizers 1 week before seed sowing. A mixture of 791 g black soil (Plantation Iwamoto Co., Ltd.), 997 g red soil (Nafco Co., Ltd.), and 259 g humus (E.H Green Corporation) per pot was sieved through an 8 mm stainless steel sieve. Approximately 500 g of mixed soil was placed in each 0.7 L pot (130 mm in diameter and 113 mm in height). During cultivation, 1 mL of 5,000-fold diluted FBP was applied weekly. Agronomic variables, including leaf number, fresh weight of leaves and shoots, and fresh biomass, were recorded at harvest.

### 2.5. Komatsuna cultivation

Komatsuna (*B. rapa*) cultivation was conducted in a greenhouse for 28 days from August to September 2023 with 50 replicates at an agricultural field in Hiroshima Prefecture, Japan. Cultivation was performed by an agricultural farmer. The treatments consisted of water (WAT) or 10,000-fold diluted FBP (FBP) application with chemical fertilizers applied according to the instructions of a fertilizer company. The soil consisted of 69.7% sand, 23.7% silt, and 6.6% clay and was classified as sandy loam. In this agricultural field, komatsuna was cultivated approximately seven times per year, and reductive soil disinfestation was performed from March to April each year. During cultivation, 10,000-fold diluted FBP was applied weekly. Agronomic variables, including fresh weights of aboveground tissues and roots and aboveground height, were recorded at harvest.

### 2.6. Analysis of bacterial community structure and bacterial copy numbers

Bacterial community structure and bacterial copy numbers were analyzed as described previously [9,10]. The MiSeq platform (Illumina, San Diego, CA, USA) was used to analyze microbial community structure. A two-stage PCR method was used for amplicon sequencing. In the first-stage PCR, the bacterial V4 region was targeted using universal primer sets, as described previously. The primer sequences were 1-515F (5′-TCG TCG GCA GCG TCA GAT GTG TAT AAG AGA CAG GTG CCA GCM GCC GCG GTA A-3′) and 1-806R (5′-GTC TCG TGG GCT CGG AGA TGT GTA TAA GAG ACA GGG ACT ACH VGG GTW TCT AAT-3′). In the second-stage PCR, purified first-stage PCR amplicons were amplified using a primer set containing flow-cell adapter, index, and tailed sequences: forward primer, 5′-AAT GAT ACG GCG ACC ACC GAG ATC TAC AC-index sequence-TCG TCG GCA GCG TC-3′; and reverse primer, 5′-CAA GCA GAA GAC GGC ATA CGA GAT-index sequence-GTC TCG TGG GCT CGG-3′. Finally, the purified PCR products from each sample were pooled, denatured, and subjected to sequencing on an Illumina MiSeq System (Illumina) using a MiSeq Reagent Kit v3 (300 × 2 cycles with paired ends; Illumina), according to the manufacturer’s instructions. The sequence data obtained from the MiSeq platform were subjected to bioinformatic analysis using QIIME2 [11]. For taxonomy-based analysis, representative sequences for each amplicon sequence variant (ASV) were analyzed using EzBioCloud [12] for bacterial data.

For quantification of bacterial cell copy numbers, real-time PCR was performed using the CFX Connect System (Bio-Rad Laboratories, Inc., Hercules, CA, USA) with universal primers targeting a portion of the bacterial 16S rRNA gene: 357F (5′-CCT ACG GGA GGC AGC AG-3′) and 518R (5′-ATT ACC GCG GCT GCT GG-3′) [13]. PCR amplification was performed as described previously [14]. To gain insight into the functional potential of the bacterial community, phylogenetic investigation of communities by reconstruction of unobserved states 2 (PICRUSt2) was used [15]. PICRUSt2 enables prediction of the number of functional genes responsible for phosphate solubilization. Plant-growth-promoting bacterial functional enzymes were quantified by multiplying the relative abundance of predicted genes by the copy number of each sample.

### 2.7. Assays of ammonium formation and phosphate solubilization

Soil samples derived from the soil immersion test, tomato cultivation, rice cultivation, and komatsuna cultivation described above were used to measure ammonium formation and phosphate solubilization. One gram of soil was suspended in 10 mL of sterilized saline water and mixed vigorously before the assay.

For measurement of ammonium formation, 100 μL of each soil sample suspension was inoculated into 10 mL of minimum nitrogen buffer containing 0.1 or 0.5 g/L peptone (Becton, Dickinson and Company, NJ, USA) and 5.0 g/L NaCl at an initial pH of 7.2. After inoculation, the buffer was incubated statically at 30 °C for 72 h. After incubation, the supernatant was recovered by centrifugation at 12,000 × *g* for 20 min, and the ammonium concentration in the supernatant was measured by the indophenol method using spectrophotometric determination of ammonia as indophenol [16]. For measurement of phosphate solubilization, 100 μL of each soil sample suspension was inoculated into 10 mL of Pikovskaya medium [17] containing 5.0 g/L insoluble tricalcium phosphate and incubated at 30 °C for 120 h with shaking at 175 rpm. After incubation, the supernatant was recovered by centrifugation at 12,000 × *g* for 20 min, and the soluble phosphate concentration in the supernatant was measured using Barton’s reagent [17].

## 3. RESULTS

### 3.1. Effect of FBP application on the growth of several agricultural crops

A tomato cultivation test was conducted for 112 days in a greenhouse at 25 °C using soil and pots supplemented with chemical fertilizer at 0% (zero), 50% (half), or 100% (full) of the recommended dosage (Table 2). Tomato seeds were sown, and 5,000-fold diluted FBP was applied weekly. The number of leaves increased as the chemical fertilizer dosage increased, but little difference was observed between treatments with and without FBP application. Similarly, plant height increased with increasing chemical fertilizer dosage, but only slight differences were observed between treatments with and without FBP application. The fresh weight of the aerial parts also increased with increasing chemical fertilizer dosage. Although the differences were not significant, FBP application increased the fresh weight of the aerial parts from 47.1 to 52.7 g and from 83.8 to 93.3 g under the 50% and 100% chemical fertilizer treatments, respectively. Root fresh weight also increased with increasing chemical fertilizer dosage. Notably, under the 50% chemical fertilizer treatment, FBP application tended to increase root fresh weight from 0.90 to 1.25 g. In addition, FBP application promoted fruit setting under the 100% chemical fertilizer treatment.

**Table 2.** Effect of FBP application on tomato growth after 112 days cultivation.

| Treatment | Leave number | Hight<br>(cm) | Fresh weight of aerial part<br>(g) | Fresh weight of root<br>(g) |
| --- | --- | --- | --- | --- |
| WAT_full | 11.0 ± 2.2 | 162 ± 17 | 83.8 ± 5.5 | 3.59 ± 1.93 |
| FBP_full | 11.0 ± 1.7 | 155 ± 9 | 93.3 ± 15.5 | 2.73 ± 1.61 |
| WAT_half | 6.00 ± 1.90 | 154 ± 9 | 47.1 ± 6.8 | 0.90 ± 0.2 |
| FBP_half | 5.80 ± 1.72 | 154 ± 11 | 52.7 ± 11.8 | 1.25 ± 0.83 |
| WAT_zero<sup>a)</sup> | 4.00 | 121 | 18.0 | 0.68 |
| FBP_zero<sup>b)</sup> | 4.50 | 103 | 18.3 | 0.34 |

Tomato cultivation test was performed applied with water (WAT) and 5.000-fold dilution FBP and chemical fertilizer dosages at 0% (zero), 50% (half), and 100% (full) (n = 5). Data are mean ± standard deviation (n = 5) except for WAT\_zero (n=1) and FBP\_zero (n=2). No significant differences were obtained between water and FBP applications.

Rice seeds (*Oryza sativa* L. ssp. *japonica* cv. Nipponbare) were sown in soil and pots supplemented with chemical fertilizer at 0% (zero), 50% (half), or 100% (full) of the recommended dosage. A 5,000-fold dilution of FBP was applied weekly. Plants were cultivated for 129 days in a greenhouse at 25 °C (Table 3). After 129 days of cultivation, FBP showed fertilizer-level-dependent effects on rice growth and yield. Under the 100% chemical fertilizer treatment, FBP slightly increased the fresh weight of the aerial parts from 43.1 to 45.2 g and the fresh weight of mature grains in husks from 11.6 to 12.2 g. Conversely, root fresh weight decreased from 41.9 to 37.1 g. The number of mature grains increased slightly from 329 to 334, and the maturity rate and thousand-kernel weight also increased from 88.1 to 89.8 and from 33.6 to 34.8 g, respectively. Under the 50% chemical fertilizer treatment, the effect of FBP on fresh weight of the aerial parts was negligible, with a minimal increase from 29.3 to 29.7 g. However, FBP had a pronounced effect on root fresh weight, which increased from 23.4 to 29.4 g. In contrast, the number of mature grains decreased from 208 to 158.

**Table 3.** Effect of FBP application on rice growth and yield after 129 days cultivation.

| Treatment | WAT_full | FBP_full | WAT_half | FBP_half | WAT_zero |
| --- | --- | --- | --- | --- | --- |
| Fresh weight of aerial part (g) | 43.1 ± 1.7 | 45.2 ± 3.2 | 29.3 ± 1.5 | 29.7 ± 2.9 | 25.1 |
| Fresh weight of root (g) | 41.9 ± 6.9 | 37.1 ± 1.9 | 23.4 ± 3.1 | 29.4 ± 2.2 | 16.4 |
| Number of mature grains | 329 ± 23 | 334 ± 24 | 208 ± 14 | 158 ± 32 | 110 |
| Number of immature grains | 45.0 ± 11.3 | 39.3 ± 30.3 | 24.7 ± 17.8 | 22.7 ± 7.6 | 9 |
| Maturity rate of grains | 88.1 ± 2.2 | 89.8 ± 7.3 | 89.6 ± 7.4 | 86.7 ± 5.9 | 92.4 |
| Fresh weight of mature grain in husks (g) | 11.6 ± 0.6 | 12.2 ± 0.8 | 7.27 ± 0.45 | 7.25 ± 0.84 | 3.54 |
| Fresh weight of immature grain in husks (g) | 0.827 ± 0.274 | 0.680 ± 0.606 | 0.497 ± 0.509 | 0.354 ± 0.303 | 0.05 |
| Dry weight of mature grain in husks (g) | 9.06 ± 0.43 | 9.49 ± 0.58 | 5.68 ± 0.32 | 5.94 ± 0.69 | 2.78 |
| Dry weight of immature grain in husks (g) | 0.630 ± 0.214 | 0.493 ± 0.415 | 0.363 ± 0.368 | 0.290 ± 0.248 | 0.03 |
| Thousand-kernel weight (g) | 33.6 ± 1.6 | 34.8 ± 1.5 | 35.4 ± 2.5 | 34.1 ± 11.8 | 34.6 |

Rice cultivation test was performed applied with water (WAT) and 5.000-fold diluted FBP and chemical fertilizer dosages at 0% (zero), 50% (half), and 100% (full) (n = 5). Data are mean ± standard deviation (n = 3) except for WAT\_zero (n=1). No significant differences were obtained between water and FBP applications.

Komatsuna seeds (*B. rapa* var. *perviridis*) were sown in soil and fields supplemented with 100% chemical fertilizer (Fig. S1). A 10,000-fold dilution of FBP was applied weekly, and a one-month cultivation trial was conducted in a greenhouse. FBP application significantly increased the height of the aerial parts from 32.8 to 34.2 cm (Fig. 1a). Although the differences were not significant, aerial part fresh weight and root length increased from 45.3 to 47.6 g and from 17.7 to 19.8 cm, respectively, with FBP application (Fig. 1b, c).

**Fig. 1.**
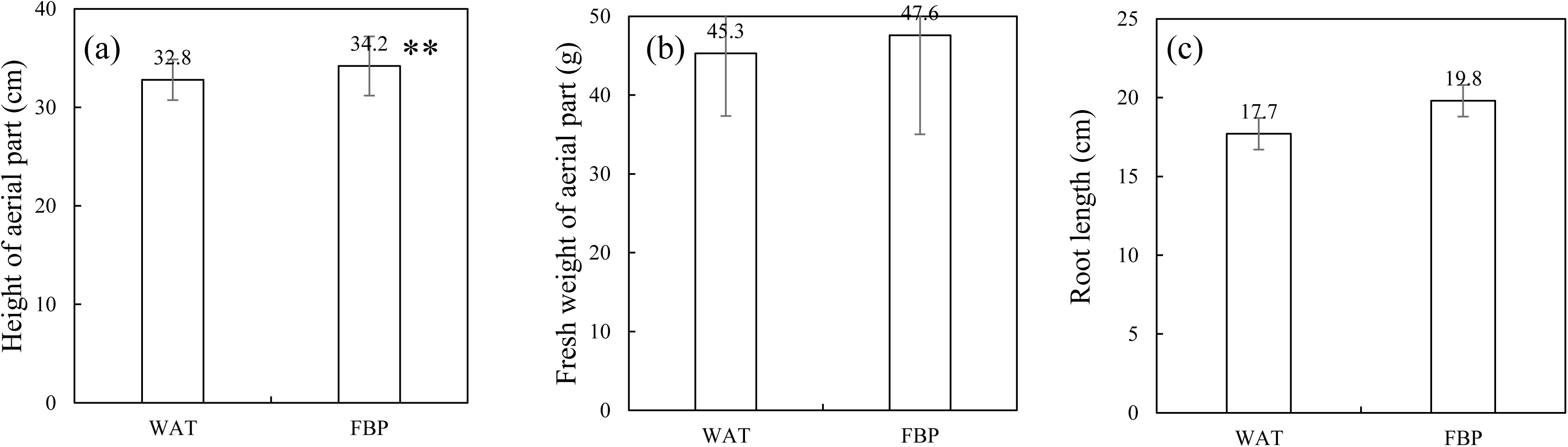
Effect of FBP application on komatsuna (*Brassica rapa*) growth after 28 days of cultivation with 10,000-fold diluted FBP or water (WAT). (a) Height of the aerial part, (b) fresh weight of the aerial part, and (c) root length. Data are presented as the mean ± standard error (n = 50). Asterisks indicate significant differences from WAT, as assessed by Student’s *t*-test: \*\**p* < 0.01.

These results indicate that FBP exerts crop- and fertilization-dependent growth-promoting effects in tomato, rice, and komatsuna.

### 3.2. Effect of FBP application on soil microbiota without crops and under several crop cultivation systems

This section analyzes bacterial communities in soils from the immersion test and tomato, rice, and komatsuna cultivation systems.

In the soil immersion test, although no significant difference was observed, application of 100-fold diluted FBP increased soil bacterial counts by approximately twofold, from 2.3 × 10⁶ to 4.5 × 10⁶ copies/g soil (Fig. S2), whereas the Shannon index decreased significantly (Fig. S3). Application of 200-fold diluted brown sugar (SUG) also significantly increased soil bacterial counts and significantly decreased the Shannon index, suggesting an effect derived from the brown sugar component of FBP. Beta-diversity analysis revealed distinct clusters corresponding to the water, brown sugar, and FBP treatments (Fig. 2). Thus, in addition to diversity changes attributable to the brown sugar component, effects of other FBP components were suggested. These results suggest that FBP application alters the soil bacterial community and bacterial counts.

**Fig. 2.**
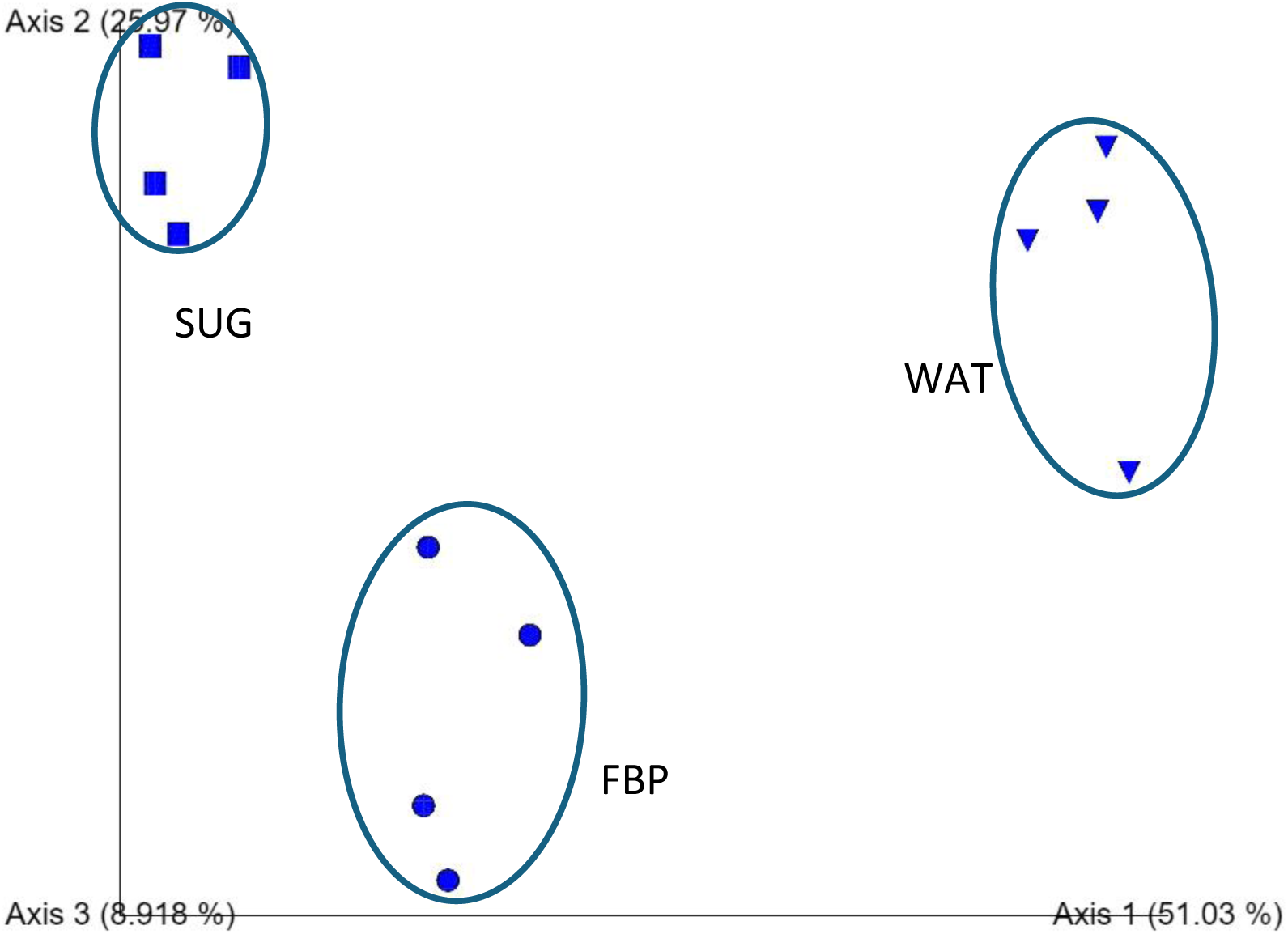
Principal coordinate analysis (PCoA) plot showing beta diversity visualized using weighted UniFrac in soil treated with water (WAT), 200-fold diluted brown sugar (SUG), or 200-fold diluted FBP for two weeks in a glass chamber (n = 4).

Soil bacterial community analysis was performed after the tomato cultivation system. Application of 5,000-fold diluted FBP did not significantly affect soil bacterial counts (Fig. S4), observed species, or the Shannon index (Table S1). Furthermore, β-diversity analysis revealed no distinct clustering among the test plots. Under the 100% and 50% chemical fertilizer treatments, FBP application significantly increased the relative abundance of the phyla Firmicutes and Spirochaetota, whereas the phylum Proteobacteria tended to decrease (Fig. S5). Furthermore, no predominant species showed significant changes in response to FBP application (Fig. S6).

Soil bacterial community analysis was performed after rice cultivation. Application of 5,000-fold diluted FBP did not significantly affect soil bacterial counts (Fig. S7), observed species, or the Shannon index (Table S2). Furthermore, β-diversity analysis revealed distinct clusters according to chemical fertilizer dosage, namely 0% (zero), 50% (half), and 100% (full), and FBP application (Fig. S8). Under the 100% and 50% chemical fertilizer treatments, FBP application increased the relative abundance of the phyla Gemmatimonadota, Desulfobacterota, and Nitrospirota (Fig. S9). Furthermore, no predominant species showed significant changes in response to FBP application (Fig. S10).

Soil bacterial community analysis was performed after komatsuna cultivation. Application of 10,000-fold diluted FBP did not significantly affect soil bacterial counts in either the rhizosphere or bulk soil, with counts maintained at approximately 10⁶ copies/g soil (Fig. S11). Furthermore, although observed species and the Shannon index did not differ significantly between treatments with and without FBP application, values tended to be higher in the rhizosphere (Table S3). Regarding β-diversity, clear clusters were formed between treatments with and without FBP application and between rhizosphere and bulk soils (Fig. 3). Under both the 100% and 50% chemical fertilizer treatments, FBP application increased the relative abundance of Gemmatimonadota in rhizosphere and bulk soils, Nitrospirota and Desulfobacterota in rhizosphere soil, and Halanaerobiaeota in bulk soil (Fig. S12). Conversely, FBP application decreased the relative abundance of the genera *Bacillus* and *Clostridium* while increasing that of *Planifilum* and *Caloramator* (Fig. S13).

**Fig. 3.**
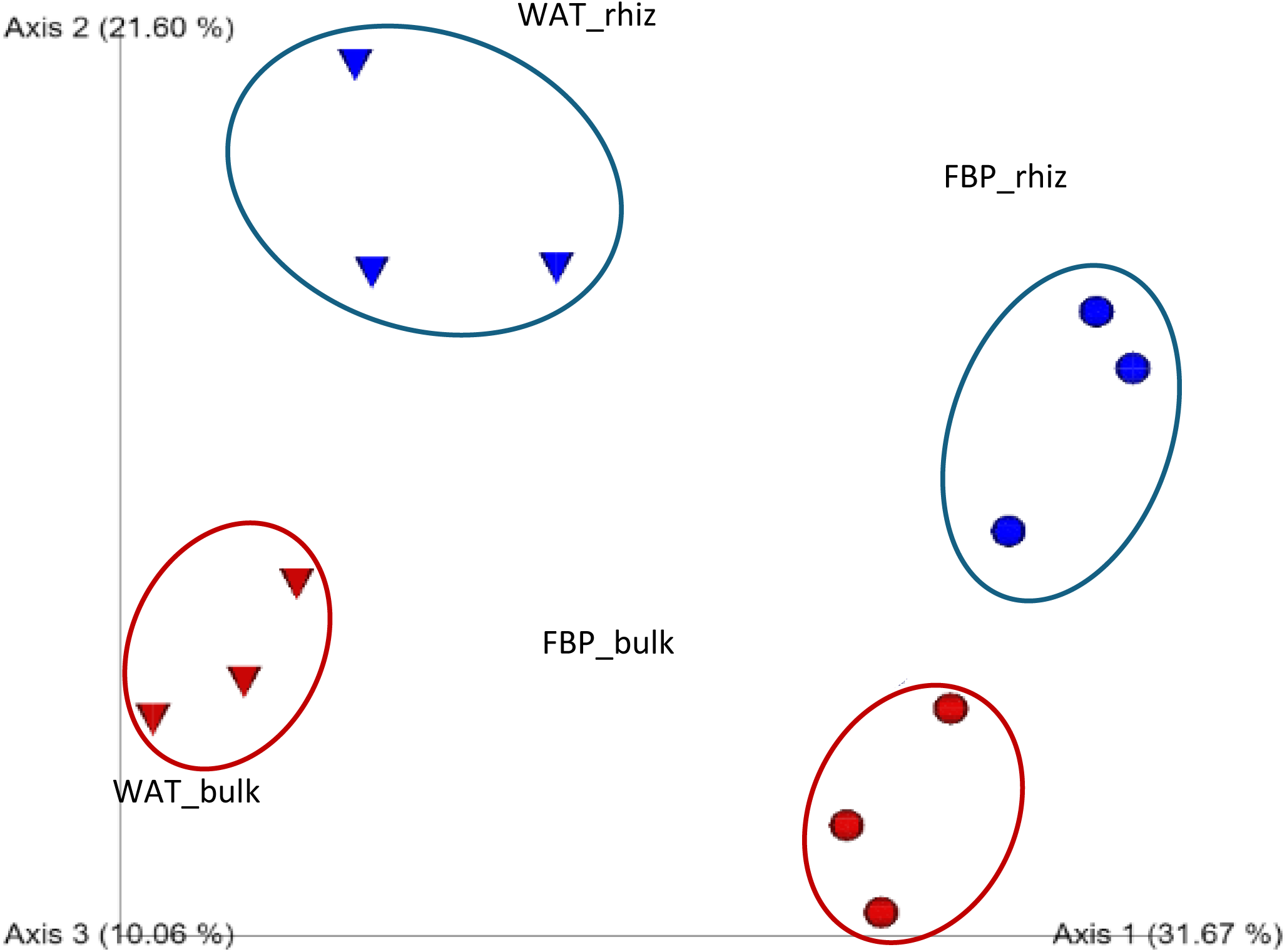
PCoA plot showing beta diversity visualized using weighted UniFrac in bulk and rhizosphere (rhiz) soils treated with 10,000-fold diluted FBP or water (WAT) after 28 days of komatsuna (*Brassica rapa*) cultivation (n = 3).

These results clearly indicate that FBP application alters the bacterial community in cultivated soil.

### 3.3. Effect of FBP application on plant-growth-promoting bacteria and ammonia formation and phosphate solubilization activities in soil microbiota under several crop cultivation systems

We measured the numbers of plant-growth-promoting bacteria in soils from the immersion test and tomato, rice, and komatsuna cultivation systems, as well as the ammonia-producing and phosphate-solubilizing activities of each soil.

For each soil, the number of plant-growth-promoting bacteria was calculated by multiplying the bacterial count by the relative abundance of bacterial species with plant growth-promoting ability. In the soil immersion test (Fig. S14), application of 100-fold diluted FBP increased *Arthrobacter methylotrophus* significantly (*p* < 0.05), as well as *Arthrobacter pokkalii*, *Arthrobacter pascens*, *Ectobacillus funiculus*, *Pseudomonas citronellolis*, and *Pseudarthrobacter niigatensis*. In tomato cultivation soil, *Paraburkholderia caledonica* increased approximately threefold following application of 5,000-fold diluted FBP, and *Caballeronia ptereochthonis* and *Paenibacillus doosanensis* were detected only with application of 5,000-fold diluted FBP. In rice cultivation soil, the number of phosphate-solubilizing *Paraburkholderia ferrariae* increased approximately fivefold following application of 5,000-fold diluted FBP compared with no water application. In komatsuna cultivation soil, application of 10,000-fold diluted FBP increased *Planifilum* sp. (Fig. S13), and *Caloramator mitchellensis* and *Sporosarcina ureilytica* were detected only in rhizosphere soils treated with 10,000-fold diluted FBP.

Soils from the immersion test and tomato, rice, and komatsuna cultivation systems were incubated at 30 °C for 3 days, and ammonia production was measured. In the soil immersion test (Fig. S15), tomato cultivation soil (Fig. S16), and rice cultivation soil (Fig. S17), FBP application did not significantly increase ammonia production. In komatsuna cultivation soil (Fig. S18 right), although the differences were not statistically significant, FBP application increased ammonia production from 23.0 to 33.4 mg/L in rhizosphere soil and from 16.7 to 21.1 mg/L in bulk soil.

Soils from the immersion test and tomato, rice, and komatsuna cultivation systems were incubated at 30 °C for 5 days, and soluble phosphate content was measured (Fig. 4). In the soil immersion test, FBP application did not significantly increase soluble phosphate content (Fig. 4a). In tomato cultivation soil, under the 0% and 100% chemical fertilizer treatments, soluble phosphate levels significantly increased with FBP application from 164 to 244 mg/L and from 131 to 244 mg/L, respectively (Fig. 4b), indicating a significant improvement in phosphate-solubilizing activity. In rice cultivation soil, under the 50% and 100% chemical fertilizer treatments, soluble phosphate levels significantly increased with FBP application from 142 to 332 mg/L and from 140 to 291 mg/L, respectively (Fig. 4c), indicating a significant improvement in phosphate-solubilizing activity. In komatsuna cultivation soil (Fig. 4d), FBP application increased soluble phosphate levels from 283 to 292 mg/L in rhizosphere soil and from 237 to 300 mg/L in bulk soil.

**Fig. 4.**
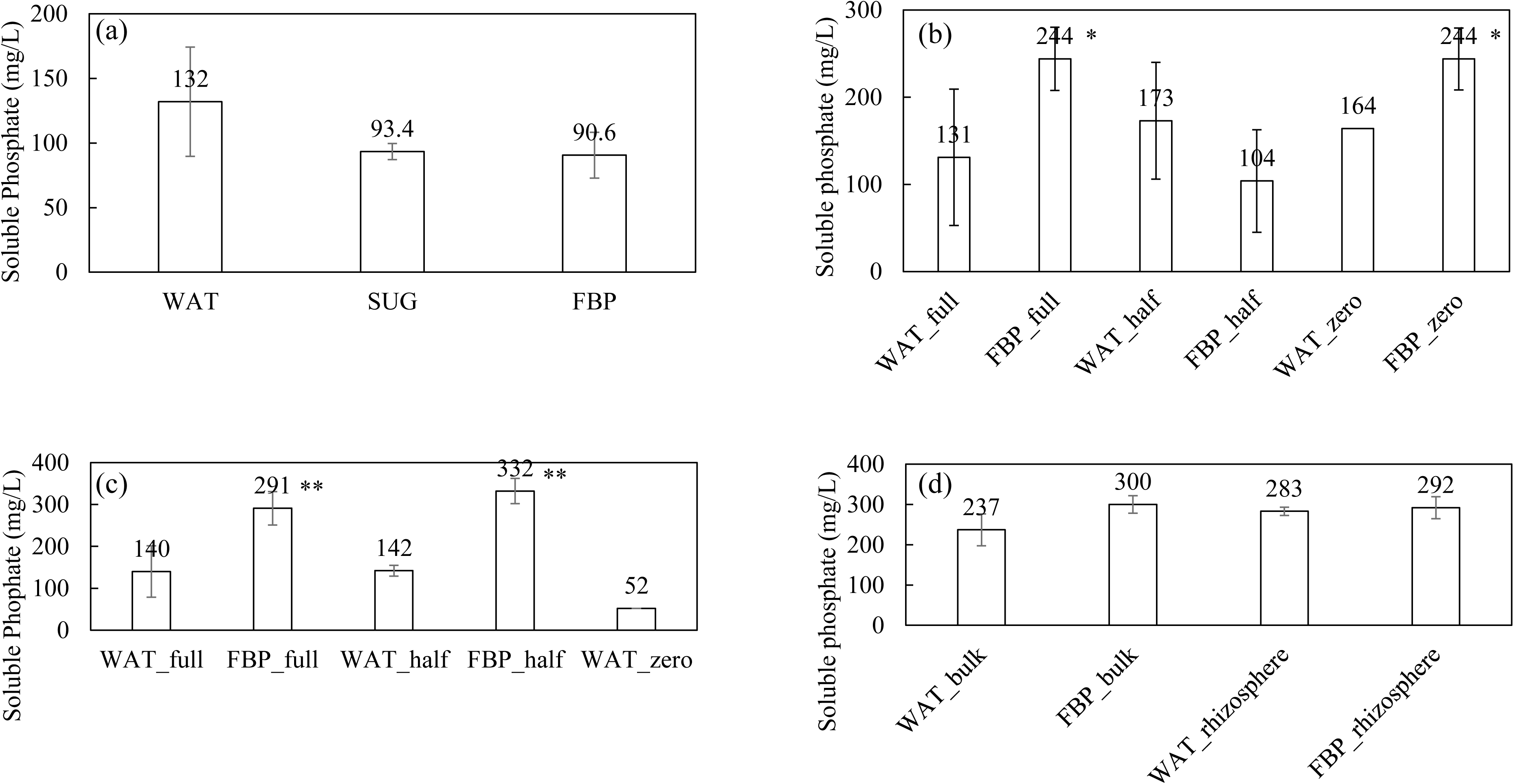
Soluble phosphate concentration in insoluble tricalcium phosphate solution after 5 days of incubation with soil-suspended solutions. (a) Soil immersion test with water (WAT), 200-fold diluted brown sugar (SUG), or 100-fold diluted FBP (n = 4); (b) tomato cultivation with WAT or 5,000-fold diluted FBP and chemical fertilizer dosages of 0% (zero), 50% (half), or 100% (full) (n = 5); (c) rice cultivation with WAT or 5,000-fold diluted FBP and chemical fertilizer dosages of zero, half, or full; and (d) bulk and rhizosphere soils from komatsuna (*Brassica rapa*) cultivation treated with water (WAT) or 10,000-fold diluted FBP (n = 3). Asterisks indicate significant differences from WAT, as assessed by Student’s *t*-test: \**p* < 0.05 and \*\**p* < 0.01.

These results suggest that FBP application alters the soil bacterial community, increases the abundance of several plant-growth-promoting bacteria, and enhances phosphate-solubilizing activity.

## 4. DISCUSSION

Achieving the SDGs requires reductions in the use of chemical fertilizers and pesticides. Interest is increasing in the adoption of biostimulants that show efficacy even under low-fertility conditions. In this study, we applied FBP to tomato, rice, and komatsuna and evaluated growth indices, soil microbial communities, and nutrient transformation functions, specifically phosphate solubilization and ammonia production. As amplification using universal primers was not achieved with DNA extracted from the FBP used in this study, FBP may exert its effects by modulating the soil microbial community and its functions rather than by directly supplying detectable live microorganisms. This section discusses crop-specific differences, the relationship between microbial communities and functions, mechanisms of action, and remaining challenges. The growth-promoting effects of plant-growth-promoting bacteria, a type of biostimulant, have previously been reported to vary depending on the cultivated variety [18]. Although growth-promoting effects of fertilization with plant-based fermented materials have been reported, few studies have examined the effects of biostimulants on the growth of different crop types (Table 4). It is also notable that this study evaluated not only the growth of several crop types but also bacterial community structures and nutrient transformation functions in the soil. In this study, the growth-promoting effects of FBP application were investigated using three crops, tomato, rice, and komatsuna (Tables 2 and 3, Fig. 1). Although the factors driving growth promotion varied by crop type, a particularly marked increase in fresh weight was observed in komatsuna. To expand the application of FBP as a biostimulant, other crops should be tested in future studies.

**Table 4.** Effect of biostimulants derived from fermented plant on crop growth in literature.

| Biostimulant | Crop | Analyzed parameters | Results | References |
| --- | --- | --- | --- | --- |
| FBP from 41 types of fruits, grains, seaweed, and root vegetables with a brown sugar | Tomato | · Vegetative / reproductive growth<br>· Bacterial community structure in soil<br>· Bacterial copy number in soil<br>· Ammonium formation activity in soil<br>· Phosphate solubilization activity in soil | · Promotion of vegetative growth<br>· Alteration of predominant phyla and increase in plant-growth-promoting bacteria numbers<br>· Increase in phosphate solubilization activity | This study |
|  | Rice |  | · Promotion of vegetative growth<br>· Alteration of beta-diversity and predominant phyla, increase in plant-growth-promoting bacteria numbers<br>· Increase in phosphate solubilization activity |  |
|  | Komatsuna<br>(*B. rapa*) |  | · Promotion of vegetative growth<br>· Alteration of alfa-diversity, beta-diversity, predominant phyla, increase in PGPB numbers<br>· Increase in ammonium formation activity<br>· Increase in phosphate solubilization activity |  |
| Fermented waste of butterfly pea flower kombucha | Eggplant | ·Vegetative / reproductive growth | ·Promotion of vegetative and reproductive growth | [24] |
| Wood vinegar and fermented bioextracts | Tomato | ·Vegetative / reproductive growth | ·Promotion of vegetative and reproductive growth | [25] |
| Plant extract from 31 agricultural materials | Lettuce | ·Vegetative growth | ·Promotion of vegetative growth | [26] |
| Commercially marketed fermentation products | Wheat | ·Vegetative growth and nitrogen uptake<br>·Microbial biomass and activities (respiration, dehydrogenase and phosphatase) | ·Promotion of vegetative growth and N uptake<br>·Increase in microbial biomass, not increase in activities | [23] |
| Fermented Alfalfa Brown Juice | Sweet Basil | ·Vegetative growth | ·Promotion of vegetative growth | [27] |
| Fermented Extract of microalgae (*Sargassum* spp.) by *Aspergillus niger* | Tomato | ·Vegetative / reproductive growth | ·Promotion of vegetative and reproductive growth | [28] |
| Fermented Kiwifruit byproduct | Wild strawberry | ·Vegetative / reproductive growth | ·Promotion of vegetative and reproductive growth | [29] |
|  | Musk strawberry |  | ·Promotion of vegetative and reproductive growth |  |
| Fermented seaweed extracts | Date palm | ·Vegetative growth<br>·Components in plant | ·Promotion of vegetative growth<br>·Increase in soluble sugar, polyphenol, peroxidase, protein | [30] |

As FBP is applied at high dilution rates of 5,000- to 10,000-fold, its direct contribution as a plant nutrient source containing nitrogen, phosphorus, and potassium is considered extremely small (Table 1). Furthermore, no 16S rRNA gene amplicons were obtained using DNA extracted from FBP, suggesting that active microorganisms, including plant-growth-promoting bacteria, do not survive in FBP. Previous studies have also reported changes in soil microbial communities and increases in plant-growth-promoting bacteria following the application of other biostimulants [10]. This study revealed that FBP application significantly alters bacterial communities regardless of crop type (Figs. 2, 3, and S8). Moreover, although the species differed by crop type, FBP application increased *A. methylotrophus*, *A. pokkalii*, *A. pascens*, *E. funiculus*, *P. citronellolis*, and *P. niigatensis* in soil-soaked samples; *P. caledonica*, *C. ptereochthonis*, and *P. doosanensis* in tomato soil; *P. ferrariae* in rice soil; and *Planifilum* sp., *C. mitchellensis*, and *S. ureilytica* in komatsuna soil (Figs. S6, S10, S13, and S14). *A. pokkalii*, *A. pascens*, and *P. citronellolis* have been reported as plant-growth-promoting bacteria that produce indole-3-acetic acid (IAA) [19–21]. Functional analysis of the soil microbiota showed that the predicted copy numbers of several genes related to phosphate solubilization increased significantly (Fig. S19). Therefore, an indirect growth-promotion mechanism mediated by FBP, associated with changes in the soil bacterial community and increases in plant-growth-promoting bacteria, is suggested. As the microbial species that increased differed according to the presence or absence of plants in the microbial community analysis, root acids secreted by plants may also influence changes in the soil microbial community. Lactic acid temporarily decreases the soil diversity index, but plant-growth-promoting bacteria remain [22]. FBP has also been confirmed to exert significant growth-promoting effects in komatsuna under sterile cultivation, as indicated by increases in leaf number, aboveground fresh weight of the aerial part, and root length (Fig. S20). Further studies are required to analyze changes in root acids resulting from direct stimulation of plants by FBP and to elucidate the mechanisms underlying its multifaceted growth-promoting effects.

The increase in plant growth-promoting activity may be attributable to the increase in the abundance of plant-growth-promoting bacteria described above. Indeed, increased plant growth-promoting activity in soil following biostimulant application has been reported. Chen et al. reported increases in β-glucosidase, phosphatase, and dehydrogenase activities following the application of an okara (soybean pulp)-fermented biostimulant [23]. However, the microbial species contributing to enhanced enzymatic activities remain unclear because soil microbial community analysis was not performed. In this study, FBP application increased ammonia-producing activity in komatsuna cultivation soil (Fig. S18) and increased phosphate-solubilizing activity in tomato, rice, and komatsuna cultivation soils (Fig. 4). Although the specific bacterial species involved remain unknown, the results suggest contributions from plant-growth-promoting bacteria, including bacterial species whose abundance increased in FBP-amended soils, as described above.

In summary, this study showed that FBP application promotes plant growth in three crops: tomato, rice, and komatsuna. The proposed mechanisms underlying FBP-mediated plant growth promotion are as follows: 1) increased numbers of plant-growth-promoting bacteria in the soil, 2) enhanced plant growth-promoting activity in the soil, and 3) direct stimulation of plants. These findings represent the first report demonstrating the suitability of FBP as a biostimulant. This study contributes to the promotion of organic agriculture, the establishment of environmentally sustainable agricultural practices, and the development of approaches to address issues such as abnormal weather conditions.

## ACKNOWLEDGEMENTS

The authors would like to acknowledge Center for Advanced Instrumental and Educational Supports, Faculty of Agriculture, Kyushu University for their support to use MiSeq and the Environmental Control Center for Experimental Biology, Kyushu University for soil immersion test and crop cultivation tests.

## CRediT authorship contribution statement

**Yukina Adachi-Oshima:** Conceptualization, Data curation, Formal analysis, Methodology, Investigation. **Ayano Hojo:** Data curation, Investigation, Formal analysis. **Yuri Mizuno:** Data curation, Investigation, Formal analysis, Writing – review and editing. **Yusuke Tateuchi:** Investigation, Supervision. **Kotaro Fujioka:** Investigation, Supervision, Writing – review and editing. **Hideto Torii:** Investigation, Supervision, Writing – review and editing. Y**ukihiro Tashiro:** Supervision. Writing – original draft, Writing – review and editing.

## Appendix A. Supplementary data

Supplementary data to this article can be found online at https: …

## Data availability

Data will be made available on request

## REFERENCES

[1] K. Chojnacka, D. Skrzypczak, D. Szopa, G. Izydorczyk, K. Moustakas, A. Witek-Krowiak, Management of biological sewage sludge: Fertilizer nitrogen recovery as the solution to fertilizer crisis, J. Environ. Manage. 326 (2023) 116602. 10.1016/j.jenvman.2022.116602.

[2] L.T.T. Liem, Y. Tashiro, P.V.T. Tinh, K. Sakai, Reduction in greenhouse gas emission from seedless lime cultivation using organic fertilizer in a province in Vietnam Mekong Delta region, Sustainability (Switzerland) 14 (2022) 6102. 10.3390/su14106102.

[3] L. Zhang, Z. Zhao, B. Jiang, B. Baoyin, Z. Cui, H. Wang, Q. Li, J. Cui, Effects of long-term application of nitrogen fertilizer on soil acidification and biological properties in China: a meta-analysis, Microorganisms 12 (2024) 1683. 10.3390/microorganisms12081683.

[4] T. Guillaume, A.M. Holtkamp, M. Damris, B. Brümmer, Y. Kuzyakov, Soil degradation in oil palm and rubber plantations under land resource scarcity, Agric. Ecosyst. Environ. 232 (2016) 110–118. 10.1016/j.agee.2016.07.002.

[5] J. Penuelas, F. Coello, J. Sardans, A better use of fertilizers is needed for global food security and environmental sustainability, Agric. Food Secur. 12 (2023) 5. 10.1186/s40066-023-00409-5.

[6] P. du Jardin, Plant biostimulants: Definition, concept, main categories and regulation, Sci. Hortic. 196 (2015) 3–14. 10.1016/j.scienta.2015.09.021.

[7] O.I. Yakhin, A.A. Lubyanov, I.A. Yakhin, P.H. Brown, Biostimulants in plant science: A global perspective, Front. Plant Sci. 7 (2017) 2049. 10.3389/fpls.2016.02049.

[8] A. Hida, N. Okano, C. Tadokoro, M. Fukunishi, A.A. Ahmed, K. Takenaka, Y. Tateuchi, K. Fujioka, H. Torii, T. Tajima, J. Kato, Fermented botanical fertilizer controls bacterial wilt of tomatoes caused by Ralstonia pseudosolanacearum, Biosci. Biotechnol. Biochem. 88 (2024) 571–576. 10.1093/bbb/zbae016.

[9] T. Koga, M. Ishizu, K. Watanabe, H. Miyamoto, M. Oshiro, K. Sakai, Y. Tashiro, Dilution rates and their transition modes influence organic acid productivity and bacterial community structure on continuous meta-fermentation using complex microorganisms, J. Biosci. Bioeng. 136 (2023) 391–399. 10.1016/j.jbiosc.2023.08.004.

[10] F. Hidayat, R.D.P. Pane, F. Sapalina, E. Listia, T. Koga, Winarna, M.E.S. Lubis, M. Oshiro, K. Sakai, S.N.H. Utami, Y. Tashiro, Long-term application of organic matter improves soil properties and plant growth-promoting bacteria in soil communities of oil palm plantation, Soil Sci. Plant Nutr. 70 (2024) 393–405. 10.1080/00380768.2024.2380881.

[11] E. Bolyen, J.R. Rideout, M.R. Dillon, N.A. Bokulich, C.C. Abnet, G.A. Al-Ghalith, H. Alexander, E.J. Alm, M. Arumugam, F. Asnicar, Y. Bai, J.E. Bisanz, K. Bittinger, A. Brejnrod, C.J. Brislawn, Reproducible, interactive, scalable and extensible microbiome data science using QIIME 2, Nat. Biotechnol. 37 (2019) 852–857. 10.1038/s41587-019-0190-3.

[12] S.H. Yoon, S.M. Ha, S. Kwon, J. Lim, Y. Kim, H. Seo, J. Chun, Introducing EzBioCloud: A taxonomically united database of 16S rRNA gene sequences and whole-genome assemblies, Int. J. Syst. Evol. Microbiol. 67 (2017) 1613–1617. 10.1099/ijsem.0.001755.

[13] K. Watanabe, A. Yamada, Y. Nishi, Y. Tashiro, K. Sakai, Host factors that shape the bacterial community structure on scalp hair shaft, Sci. Rep. 11 (2021) 17711. 10.1038/s41598-021-96767-w.

[14] M. Zhang, Y. Tashiro, Y. Asakura, N. Ishida, K. Watanabe, S. Yue, M.N. Akiko, K. Sakai, Lab-scale autothermal thermophilic aerobic digestion can maintain and remove nitrogen by controlling shear stress and oxygen supply system, J. Biosci. Bioeng. 132 (2021) 293–301. 10.1016/j.jbiosc.2021.05.008.

[15] M.G.I. Langille, J. Zaneveld, J.G. Caporaso, D. McDonald, D. Knights, J.A. Reyes, J.C. Clemente, D.E. Burkepile, R.L. Vega Thurber, R. Knight, R.G. Beiko, C. Huttenhower, Predictive functional profiling of microbial communities using 16S rRNA marker gene sequences, Nat. Biotechnol. 31 (2013) 814–821. 10.1038/nbt.2676.

[16] Y. Tashiro, K. Kanda, Y. Asakura, T. Kii, H. Cheng, P. Poudel, Y. Okugawa, K. Tashiro, K. Sakai, A unique autothermal thermophilic aerobic digestion process showing a dynamic transition of physicochemical and bacterial characteristics from the mesophilic to the thermophilic phase, Appl. Environ. Microbiol. 84 (2018) 1–15. 10.1128/AEM.02537-17.

[17] A. Pande, P. Pandey, S. Mehra, M. Singh, S. Kaushik, Phenotypic and genotypic characterization of phosphate solubilizing bacteria and their efficiency on the growth of maize, J Genet. Eng. Biotechnol. 15 (2017) 379–391. 10.1016/j.jgeb.2017.06.005.

[18] Y. Li, Y. Li, H. Zhang, M. Wang, S. Chen, Diazotrophic Paenibacillus beijingensis BJ- 18 Provides nitrogen for plant and promotes plant growth, nitrogen uptake and metabolism, Front. Microbiol. 10 (2019) 1119. 10.3389/fmicb.2019.01119.

[19] R. Krishnan, R.R. Menon, N. Tanaka, H.J. Busse, S. Krishnamurthi, N. Rameshkumar, *Arthrobacter pokkalii* sp nov, a novel plant associated actinobacterium with plant beneficial properties, isolated from saline tolerant pokkali rice, Kerala, India, PLoS One 11 (2016) e0150322. 10.1371/journal.pone.0150322.

[20] M. Li, R. Guo, F. Yu, X. Chen, H. Zhao, H. Li, J. Wu, Indole-3-acetic acid biosynthesis pathways in the plant-beneficial bacterium *Arthrobacter pascens* zz21, Int. J. Mol. Sci. 19 (2018) 443. 10.3390/ijms19020443.

[21] M.B. Sulochana, S.Y. Jayachandra, S.A. Kumar, A.B. Parameshwar, K.M. Reddy, A. Dayanand, Siderophore as a potential plant growth-promoting agent produced by *Pseudomonas aeruginosa* JAS-25, Appl. Biochem. Biotechnol. 174 (2014) 297–308. 10.1007/s12010-014-1039-3.

[22] S. Macias-Benitez, A.M. Garcia-Martinez, P. Caballero Jimenez, J.M. Gonzalez, M. Tejada Moral, J. Parrado Rubio, Rhizospheric organic acids as biostimulants: monitoring feedbacks on soil microorganisms and biochemical properties, Front. Plant Sci. 11 (2020) 633. 10.3389/fpls.2020.00633.

[23] S.-K. Chen, C.A. Edwards, S. Subler, The influence of two agricultural biostimulants on nitrogen transformations, microbial activity, and plant growth in soil microcosms, Soil Biol. Biochem. 35 (2003) 9–19. www.elsevier.com/locate/soilbio.

[24] M. Faizal Fathurrohim, F. Hidayanto, F. Rezaldi, Y. Kolo, halal biotechnology on fermentation and liquid fertilizer preparation from kombucha waste of tecablowe waste in increasing eggplant (*Solanum molengena*) growth, Int. J. Mathla’ul Anwar Halal Issues 2 (2022) 85–92.

[25] T. Mungkunkamchao, T. Kesmala, S. Pimratch, B. Toomsan, D. Jothityangkoon, Wood vinegar and fermented bioextracts: Natural products to enhance growth and yield of tomato (*Solanum lycopersicum* L.), Sci. Hortic. 154 (2013) 66–72. 10.1016/j.scienta.2013.02.020.

[26] S.J. Jang, Y.I. Kuk, Growth promotion effects of plant extracts on various leafy vegetable crops, Hortic. Sci. Technol. 37 (2019) 322–336. 10.7235/HORT.20190033.

[27] S. Kisvarga, D. Barna, S. Kovács, G. Csatári, I.O. Tóth, M.G. Fári, P. Makleit, S. Veres, T. Alshaal, N. Bákonyi, Fermented alfalfa brown juice significantly stimulates the growth and development of sweet basil (*Ocimum basilicum* L.) plants, Agronomy 10 (2020). 10.3390/agronomy10050657.

[28] R.M. Paredes-Camacho, A. Robledo-Olivo, S. González-Morales, A. Benavides-Mendoza, R.M. Rodríguez-Jasso, J.A. González-Fuentes, A.V. Charles-Rodríguez, Evaluation of the fermented extract of *Sargassum* spp., for the biostimulation in the germination of tomato seeds and seedlings (*Solanum lycopersicum* L.), J. Soil Sci. Plant Nutr. (2024). 10.1007/s42729-024-01877-9.

[29] S. Nazeer, A. Agosti, L. Del Vecchio, L. Leto, A. Di Fazio, J.H. Saadoun, A. Levante, C. Lazzi, M. Cirlini, B. Chiancone, Assessment of fermented kiwifruit on morpho-physiological and productive performances of *Fragaria* spp plants, grown under hydroponic conditions, J. Sustain. Agric. Environ/ 3 (2024) 1–12. 10.1002/sae2.70024.

[30] A. Nogot, M. Akaddar, A. Khardi, A. El Abbassi, F. Jaiti, Enhancing young date palm *Boufeggouss* cv. seedling tolerance to drought stress with fermented seaweed extracts from Gelidium sesquipedale, EuroMediterr. J. Environ. Integr. 9 (2024) 2123–2135. 10.1007/s41207-024-00574-4.

