## Supplemental Table for "Fermented Botanical Product Modulates Soil Bacterial Communities and Enhances Plant-Growth-Promoting Activity for Sustainable Agriculture"

Table S1 Alfa-diversity indexes of microbiota in soil applied with FBP after 112 days of tomato cultivation.

| Treatment | observed_features | shannon_entropy | faith_pd | pielou_evenness |
|---|---|---|---|---|
| wat-full1 | 991 | 9.26 | 50.6 | 0.931 |
| wat-full2 | 888 | 9.05 | 48 | 0.924 |
| wat-full3 | 533 | 8.47 | 36.1 | 0.935 |
| wat-full4 | 430 | 8.05 | 31.2 | 0.92 |
| wat-full5 | 210 | 4.79 | 39.8 | 0.621 |
| wat-half1 | 707 | 8.83 | 44 | 0.932 |
| wat-half2 | 529 | 8.48 | 36.1 | 0.937 |
| wat-half3 | 1106 | 9.38 | 51.6 | 0.927 |
| wat-half4 | 182 | 6.92 | 21.5 | 0.922 |
| wat-half5 | 797 | 8.98 | 45.8 | 0.931 |
| fbp-full1 | 936 | 9.15 | 51.1 | 0.927 |
| fbp-full2 | 825 | 9.02 | 44.7 | 0.931 |
| fbp-full3 | 927 | 9.18 | 50.6 | 0.932 |
| fbp-full4 | 688 | 8.74 | 43.1 | 0.927 |
| fbp-full5 | 724 | 8.81 | 44.4 | 0.928 |
| fbp-half1 | 832 | 9.04 | 45.8 | 0.932 |
| fbp-half2 | 547 | 5.83 | 72.5 | 0.641 |
| fbp-half3 | 858 | 9.1 | 47.2 | 0.934 |
| fbp-half4 | 537 | 8.42 | 37.4 | 0.928 |
| fbp-half5 | 660 | 7.71 | 69.9 | 0.823 |
| wat-zero | 1033 | 9.06 | 90.7 | 0.905 |
| fbp-zero1 | 439 | 8.26 | 57.7 | 0.941 |
| fbp-zero2 | 803 | 7.22 | 51 | 0.748 |

Tomato cultivation was performed applied with water (WAT) and 5,000-diluted FBP and chemical fertilizer dosages at 0% (zero), 50% (half), and 100% (full).

Table S2 Alfa-diversity indexes of microbiota in soil applied with FBP after 129 days of rice cultivation.

| Treatment | observed_features | shannon_entropy | faith_pd | pielou_evenness |
|---|---|---|---|---|
| WAT_full1 | 685 | 8.46 | 38.0 | 0.898 |
| WAT_full2 | 782 | 8.93 | 43.6 | 0.929 |
| WAT_full3 | 588 | 8.42 | 35.2 | 0.916 |
| WAT_full4 | 625 | 8.41 | 34.8 | 0.906 |
| FBP_full1 | 588 | 8.44 | 36.7 | 0.917 |
| FBP_full2 | 544 | 8.44 | 42.2 | 0.929 |
| FBP_full3 | 572 | 8.32 | 35.8 | 0.908 |
| WAT_half1 | 631 | 8.42 | 36.1 | 0.905 |
| WAT_half2 | 497 | 8.16 | 32.5 | 0.911 |
| WAT_half3 | 741 | 8.77 | 50.2 | 0.920 |
| FBP_half1 | 579 | 8.26 | 36.1 | 0.900 |
| FBP_half2 | 367 | 7.78 | 28.5 | 0.913 |
| FBP_half3 | 544 | 8.34 | 36.0 | 0.917 |
| WAT_zero | 400 | 8.02 | 28.9 | 0.927 |

Rice cultivation was performed applied with water (WAT) and 5,000-diluted FBP and chemical fertilizer dosages at 0% (zero), 50% (half), and 100% (full).

Table S3 Alfa-diversity indexes of microbiota in bulk and rhizosphere soils applied with FBP after 28 days of komatsuna (*B. rapa)* cultivation.

| Treatments | Soil | observed_features | shannon_entropy | faith_pd | pielou_evenness |
|---|---|---|---|---|---|
| Water | Bulk | 750 | 8.86 | 58.9 | 0.928 |
| Water | Bulk | 558 | 8.51 | 46.6 | 0.933 |
| Water | Bulk | 842 | 8.9 | 64.7 | 0.916 |
| Water | Rhizosphere | 958 | 9.15 | 71.1 | 0.924 |
| Water | Rhizosphere | 650 | 8.77 | 54.4 | 0.939 |
| Water | Rhizosphere | 1050 | 9.26 | 73.8 | 0.922 |
| FBP | Bulk | 626 | 8.56 | 52.5 | 0.922 |
| FBP | Bulk | 513 | 8.33 | 45.7 | 0.925 |
| FBP | Bulk | 1010 | 9.13 | 66 | 0.915 |
| FBP | Rhizosphere | 651 | 8.6 | 53.3 | 0.92 |
| FBP | Rhizosphere | 1067 | 9.25 | 73.3 | 0.92 |
| FBP | Rhizosphere | 1129 | 9.31 | 73.6 | 0.918 |

Komatsuna cultivation was performed applied with water (WAT) and 10,000-diluted FBP.
