## Supplemental Figure for "Fermented Botanical Product Modulates Soil Bacterial Communities and Enhances Plant-Growth-Promoting Activity for Sustainable Agriculture"

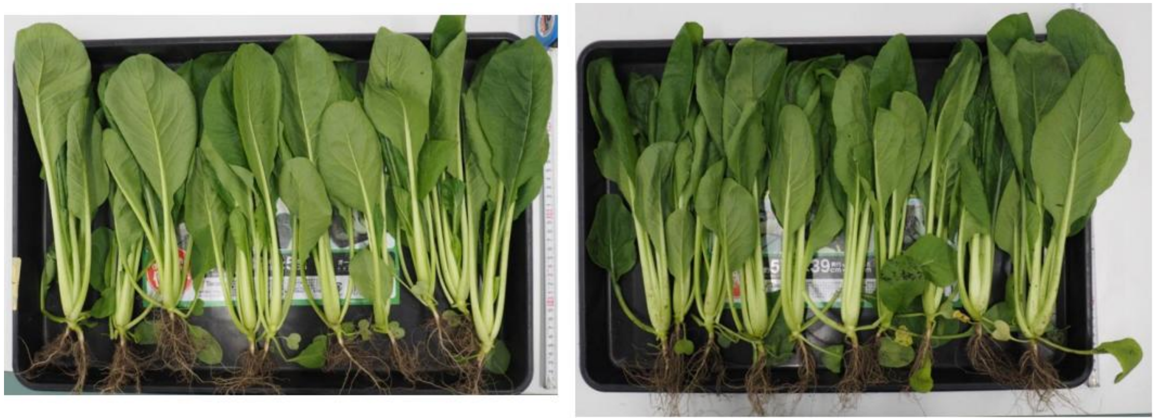

**WAT** **FBP**

Fig. S1 Photographs of komatsuna (*B. rapa)* body after 28 days of cultivation applied with 10,000-fold diluted FBP (right) or water (WAT) (left).

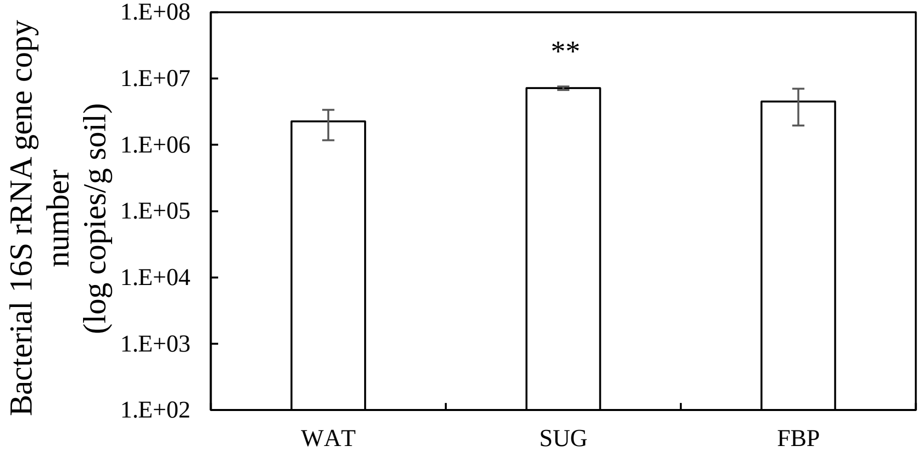


Fig. S2 Bacterial copy numbers in soil applied with water (WAT), 200-fold diluted brown sugar (SUG) or 100-fold diluted FBP for two weeks in a glass chamber. Data are presented as the mean ± standard error bar (n = 3). Asterisks indicate significant differences from WAT, as assessed by Student's t-test: **$p < 0.01$.

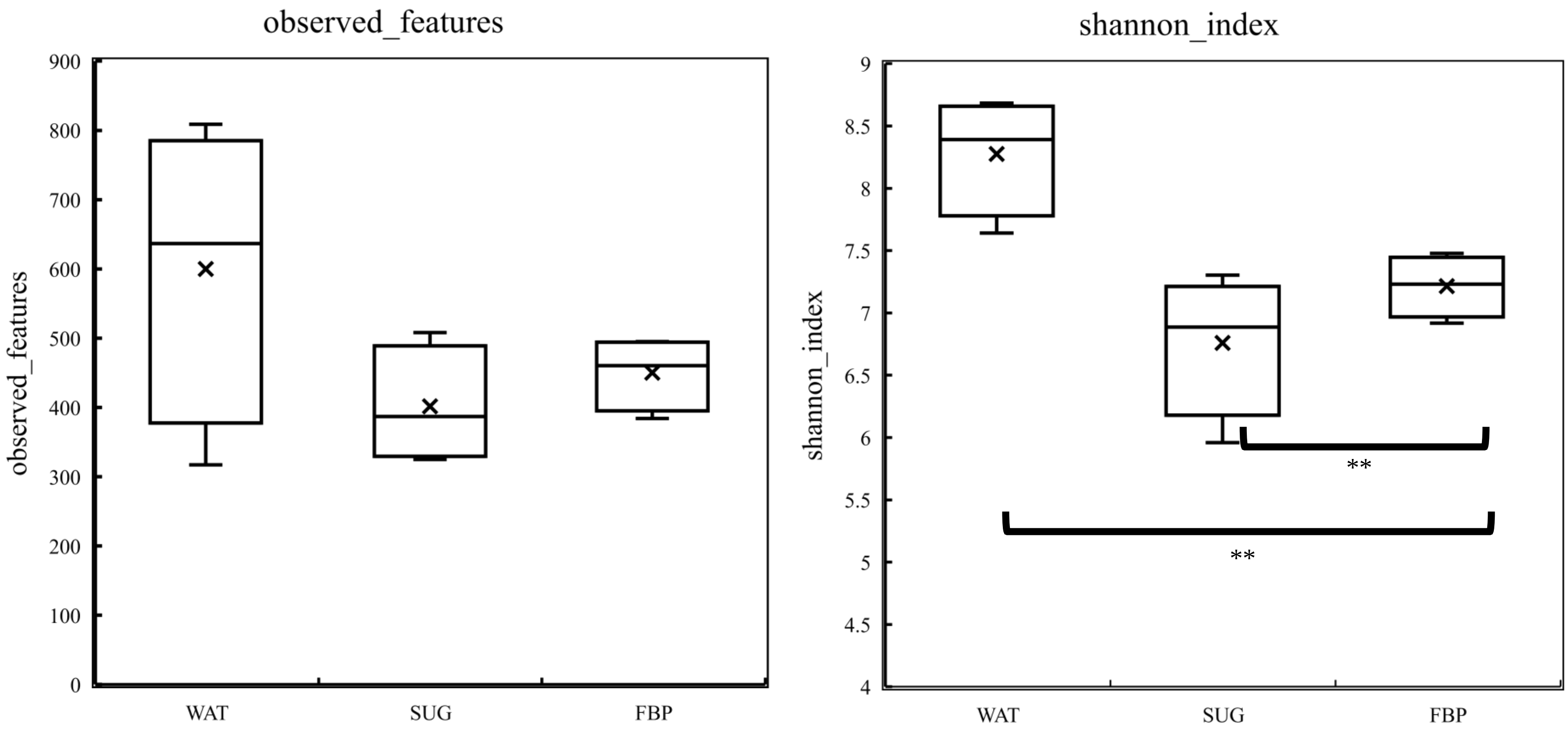


Fig. S3 Alfa-diversity indexes of observed features (left) and shannon (right) in soil applied with water (WAT), 200-fold diluted brown sugar (SUG) and 100-fold diluted FBP for two weeks in a glass chamber. Data are presented as the mean ± standard error bar (n=4). Asterisks indicate significant differences from WAT, as assessed by Student's t-test: *$p < 0.05$, **$p < 0.01$.

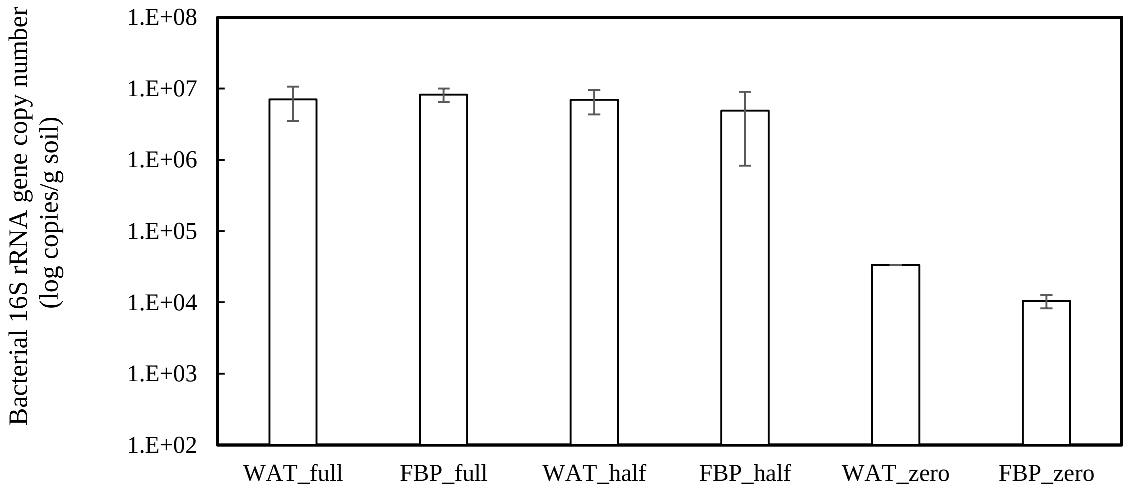


Fig. S4 Bacterial copy numbers in soil applied with 5,000-fold diluted FBP after 112 days of tomato cultivation applied with water (WAT) or 5,000-fold diluted FBP and chemical fertilizer dosages at 0% (zero), 50% (half), and 100% (full). Data are presented as the mean ± standard error bar (n=5) except for WAT_zero (n=1) and FBP_zero (n=2). The statistical method used was Student t-test. No significant differences were obtained between WAT and FBP applications.

Tomato cultivation test was performed applied with water (WAT) and FBP and chemical fertilizer dosages at 0% (zero), 50% (half), and 100% (full)

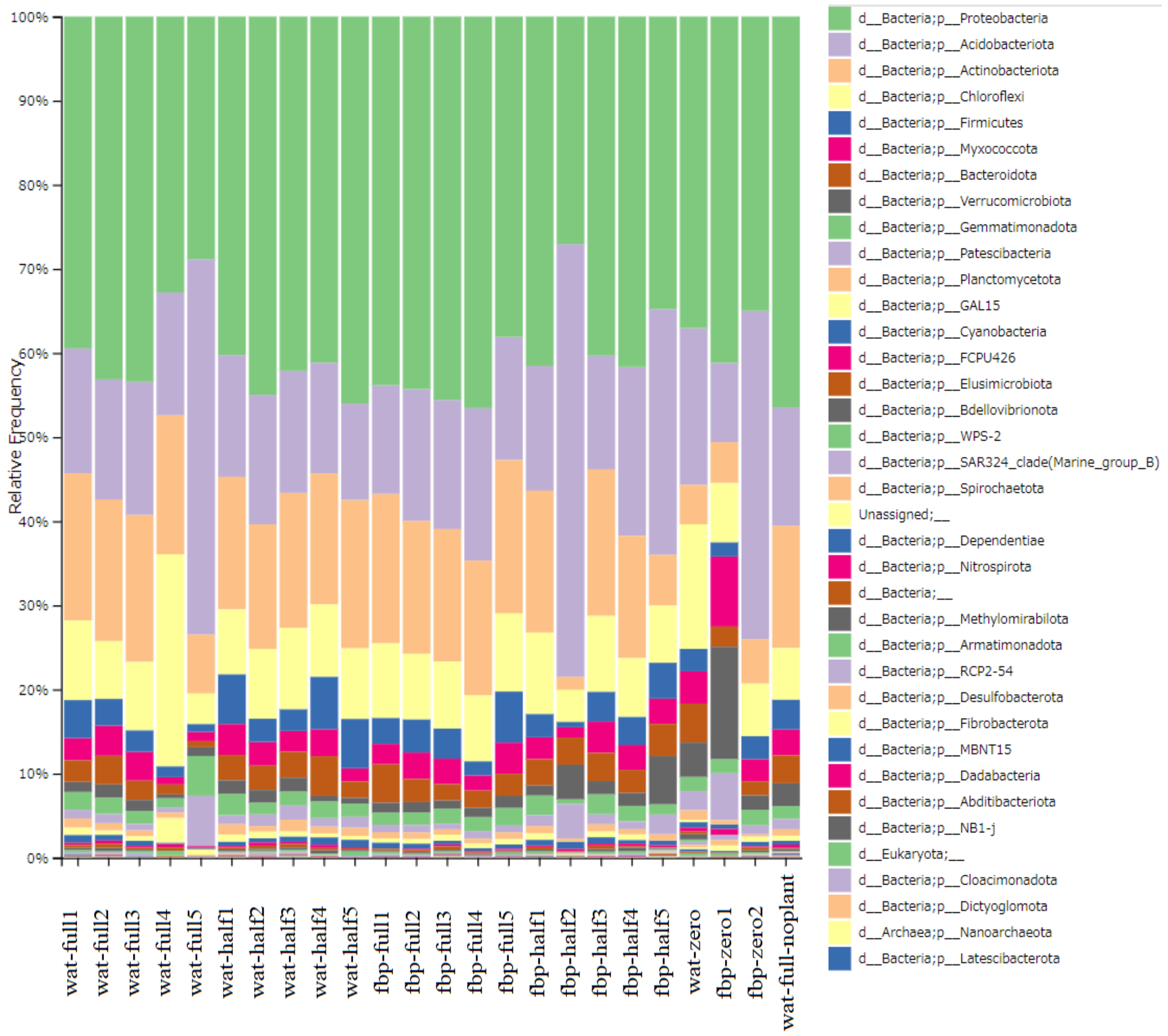


Fig. S5 Relative abundance of predominant phyla in soil applied with 5,000-fold diluted FBP after 112 days of tomato cultivation applied with water (WAT) or 5,000-fold diluted FBP and chemical fertilizer dosages at 0% (zero), 50% (half), and 100% (full).

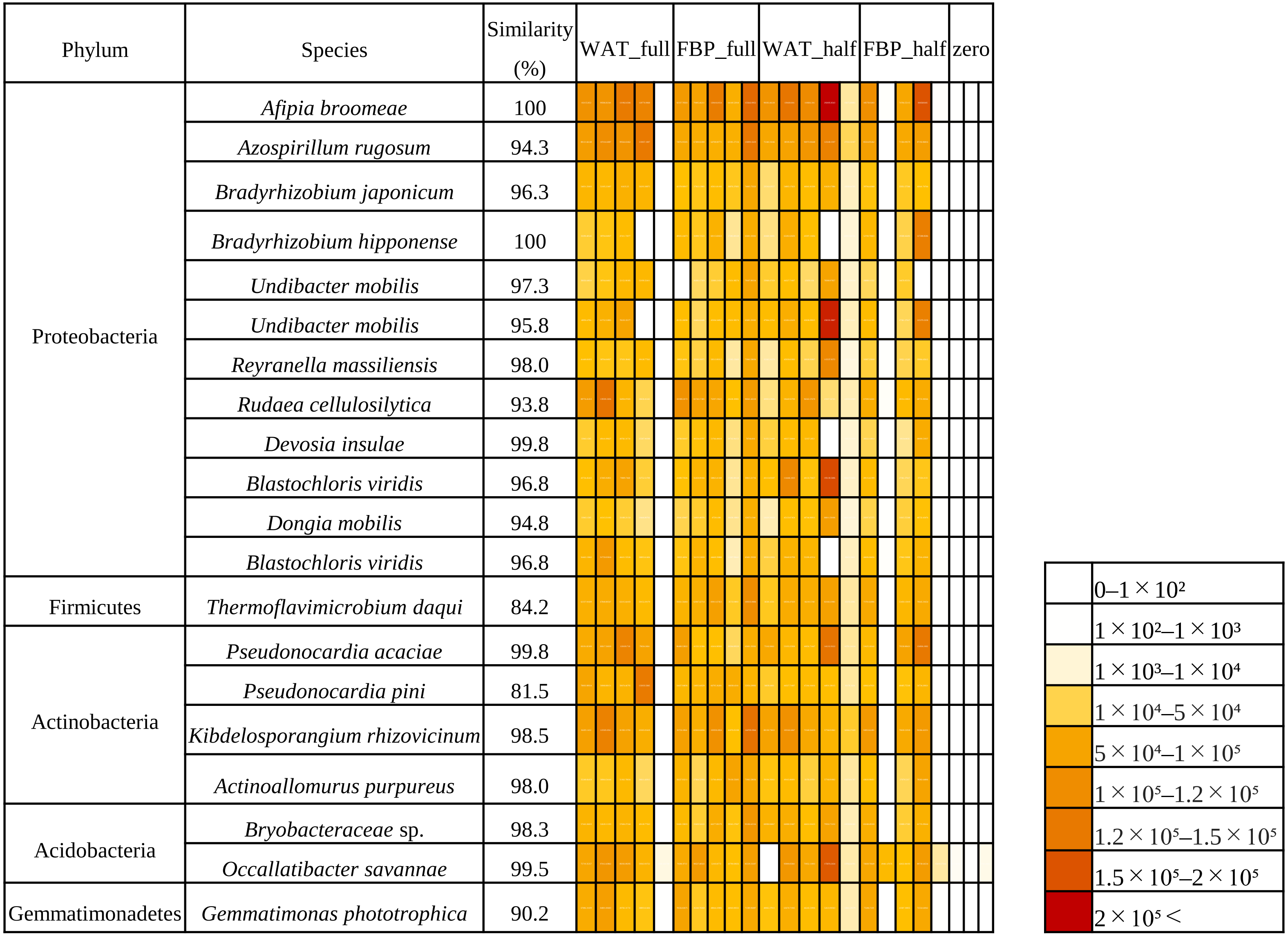


Fig. S6 Heatmap of predominant species copy number in soils after 112 days of tomato cultivation applied with water (WAT) or 5,000-fold diluted FBP and chemical fertilizer dosages at 0% (zero), 50% (half), and 100% (full)

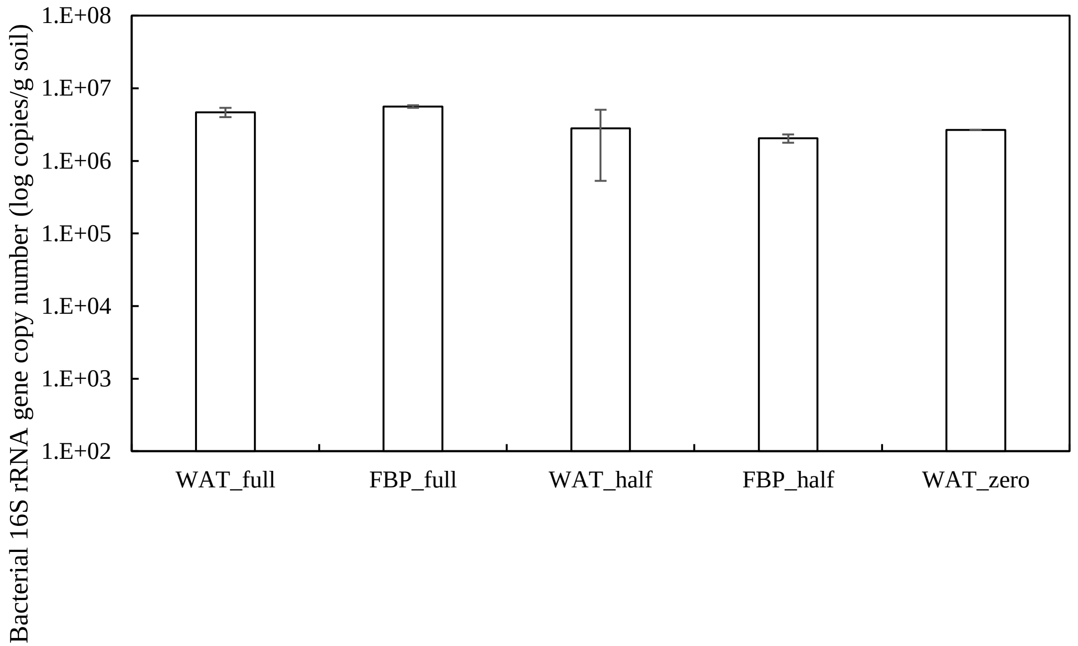


Fig. S7 Bacterial copy numbers in soil after 129 days of rice cultivation applied with water (WAT) or 5,000-fold diluted FBP and chemical fertilizer dosages at 0% (zero), 50% (half), and 100% (full). Data are presented as the mean ± standard error bar (n=3) except for WAT_zero (n=1). The statistical method used was Student t-test. No significant differences were obtained between water and FBP applications.

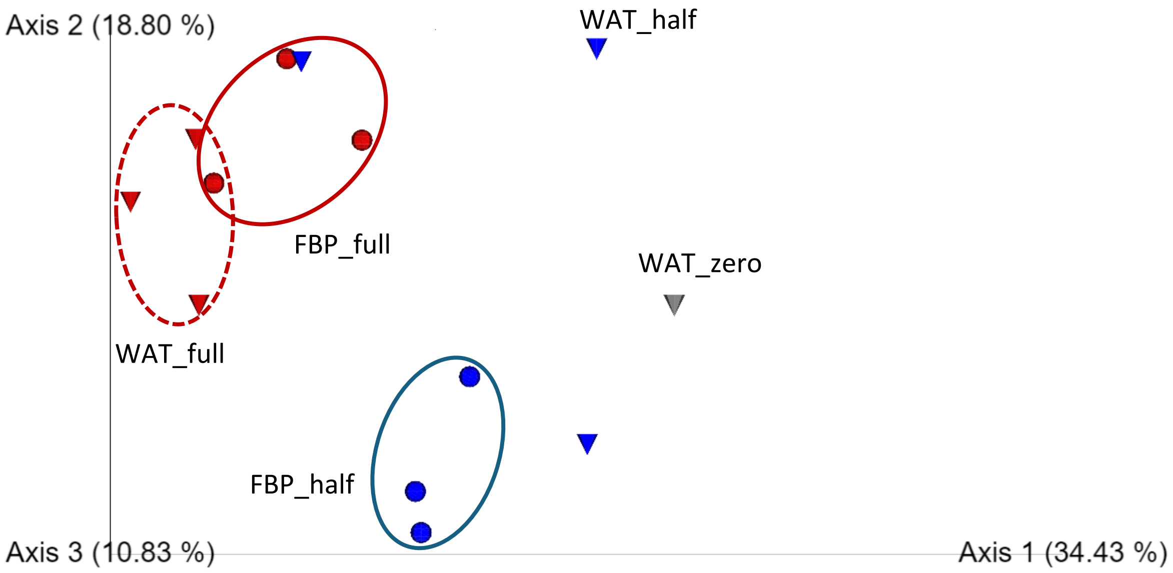


Fig. S8 Principal coordinate analysis (PCoA) plot showing beta diversity visualized using weighted-UniFrac after 129 days of rice cultivation in soil applied with water (WAT) or 5,000-fold diluted FBP and chemical fertilizer dosages at 0% (zero), 50% (half), and 100% (full).

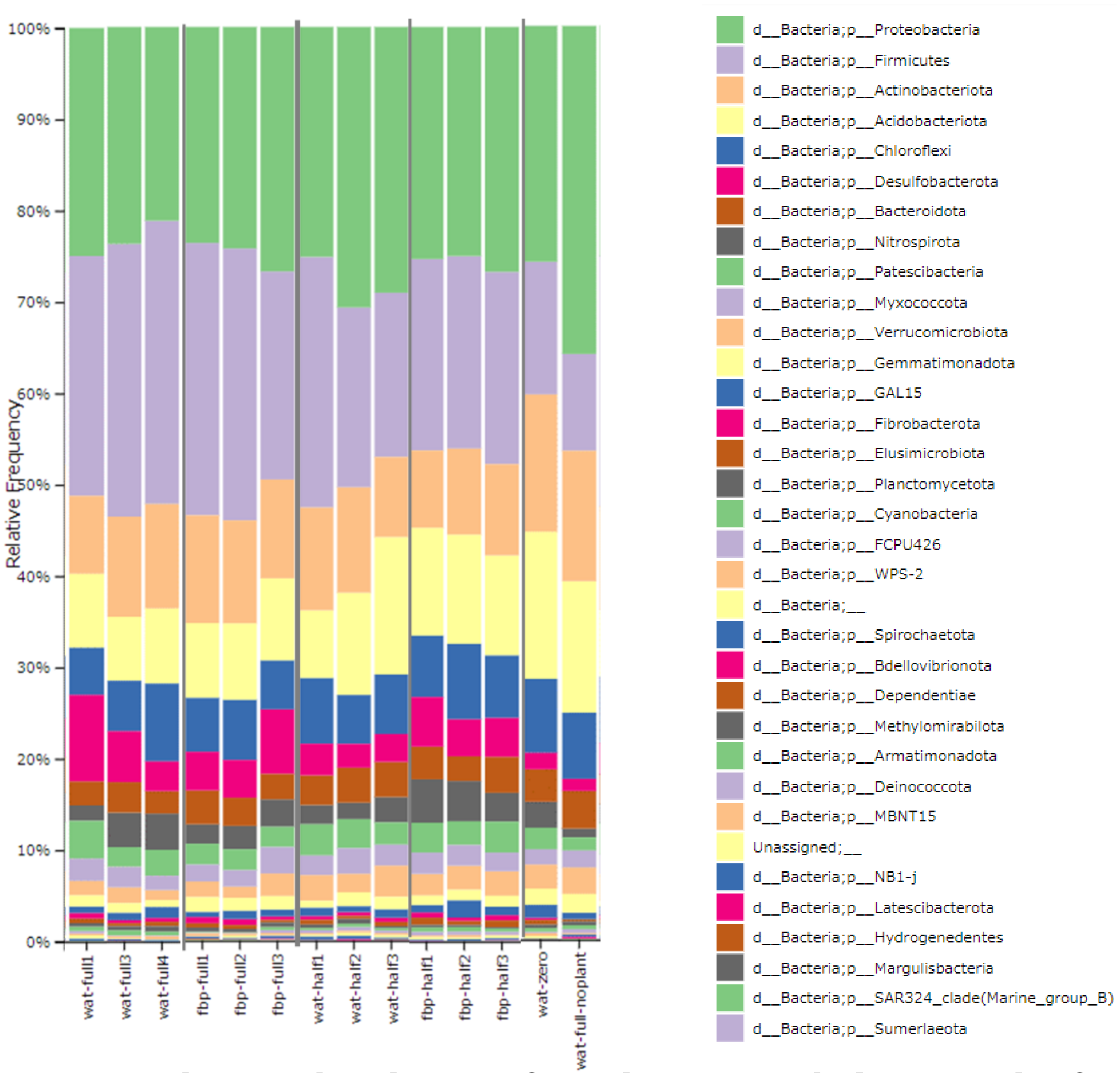


Fig. S9 Relative abundance of predominant phyla in soil after 129 days of rice cultivation applied with water (WAT) or 5,000-fold diluted FBP and chemical fertilizer dosages at 0% (zero), 50% (half), and 100% (full).

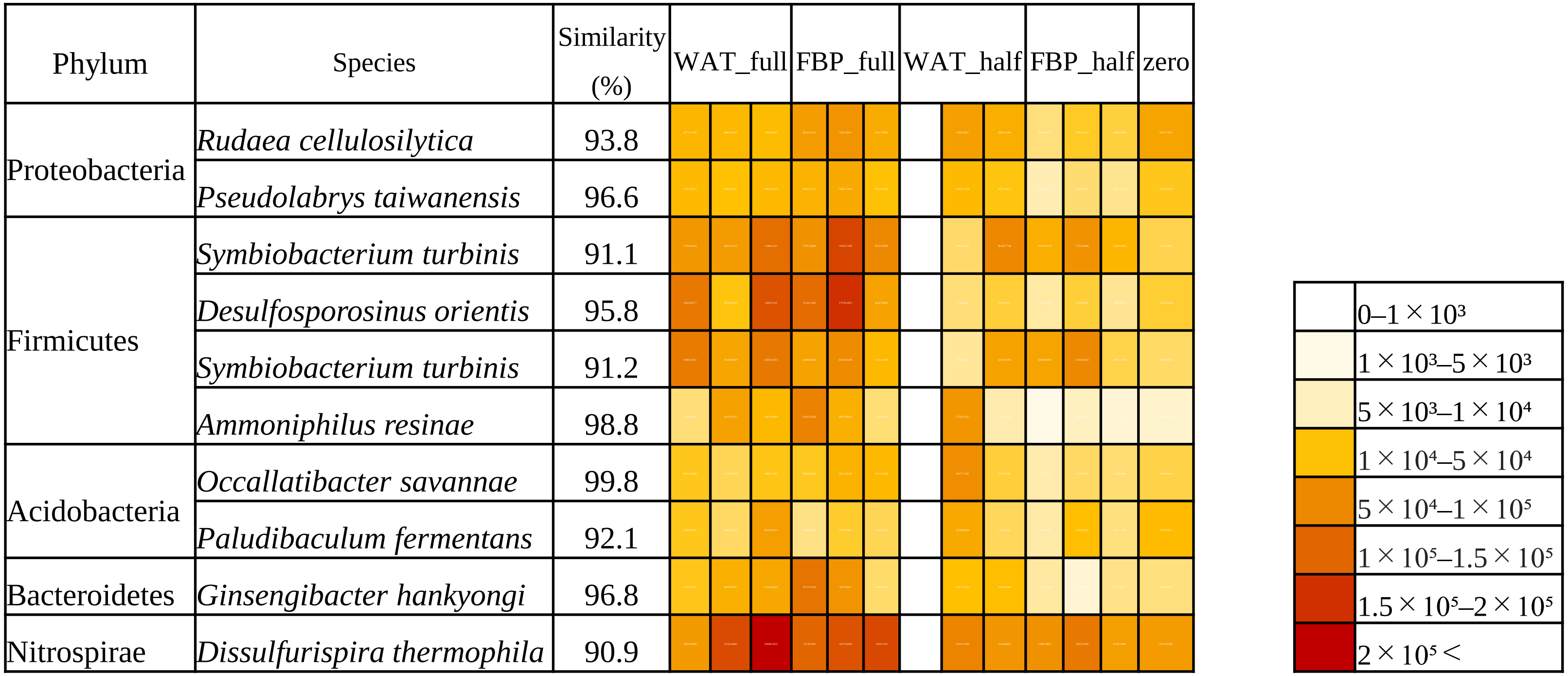


Fig. S10 Heatmap of predominant species copy number in soil after 129 days of rice cultivation applied with water (WAT) or 5,000-fold diluted FBP and chemical fertilizer dosages at 0% (zero), 50% (half), and 100% (full).

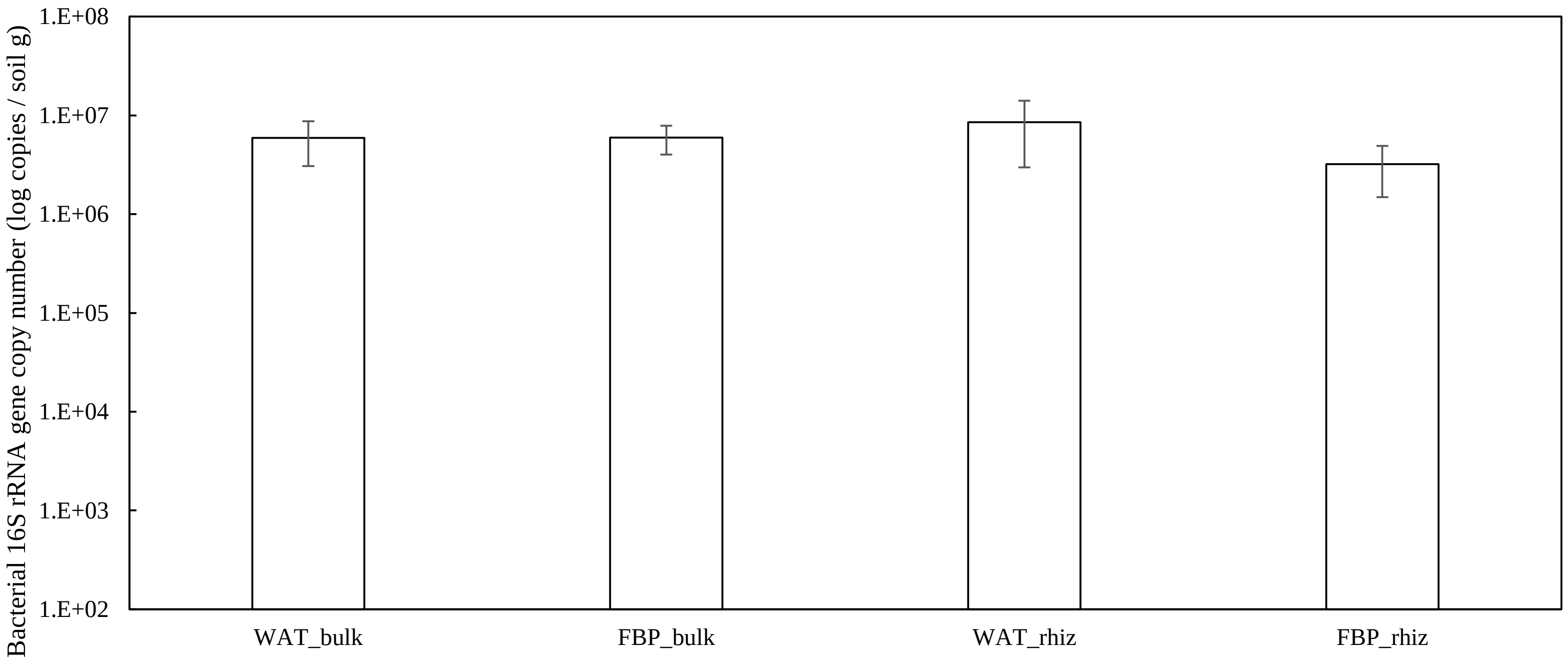


Fig. S11 Bacterial copy numbers in bulk and rhizosphere (rhiz) soils applied with water (WAT) or 10,000-fold diluted FBP after 28 days of komatsuna (*B. rapa)* cultivation. Data are presented as the mean ± standard error bar (n=3). The statistical method used was Student t-test. No significant differences were obtained between water and FBP applications.

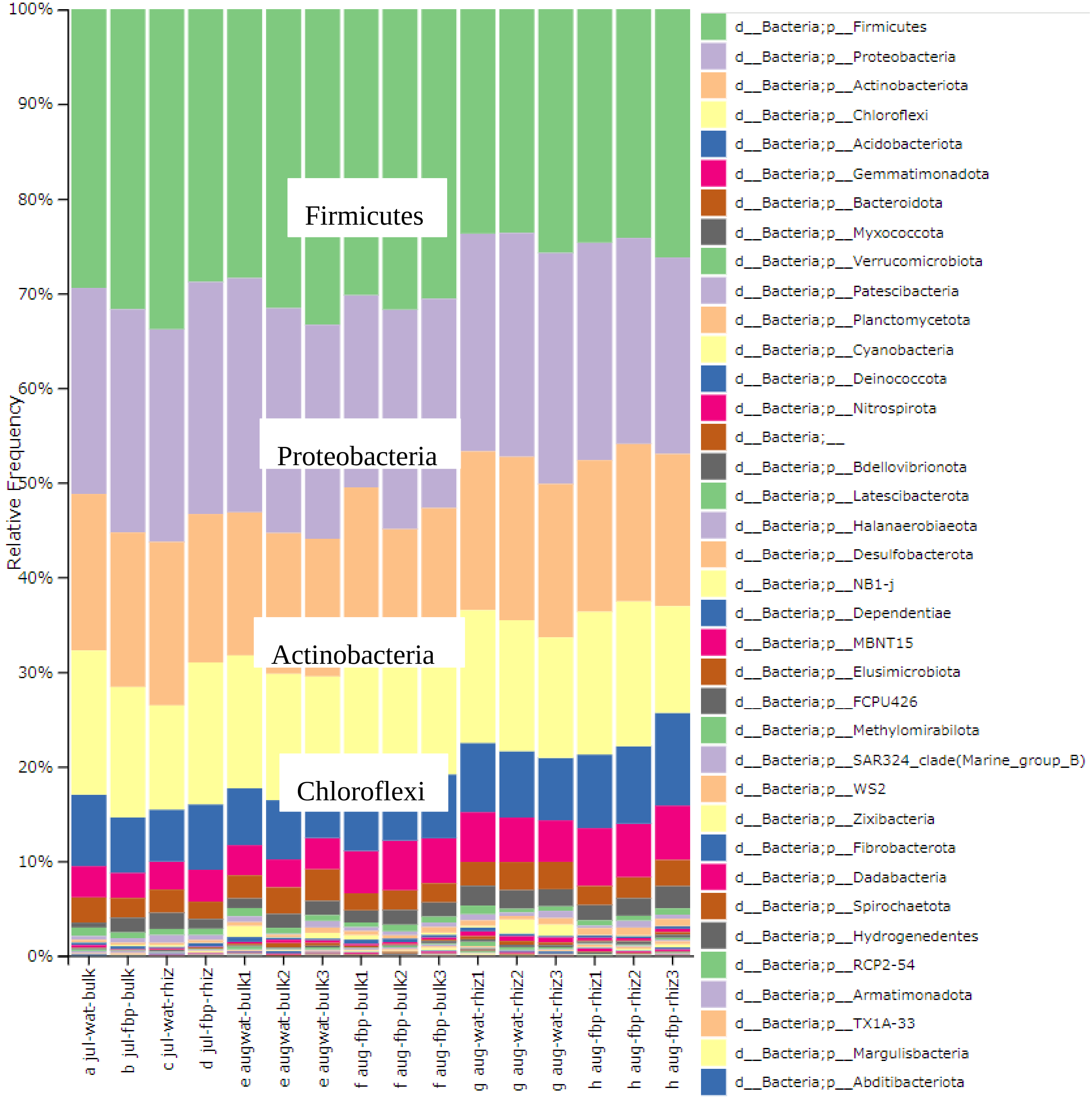


Fig. S12 Relative abundance of predominant phyla in bulk and rhizosphere soils applied with water (WAT) or 10,000-fold diluted FBP after 28 days of komatsuna (*B. rapa)* cultivation.

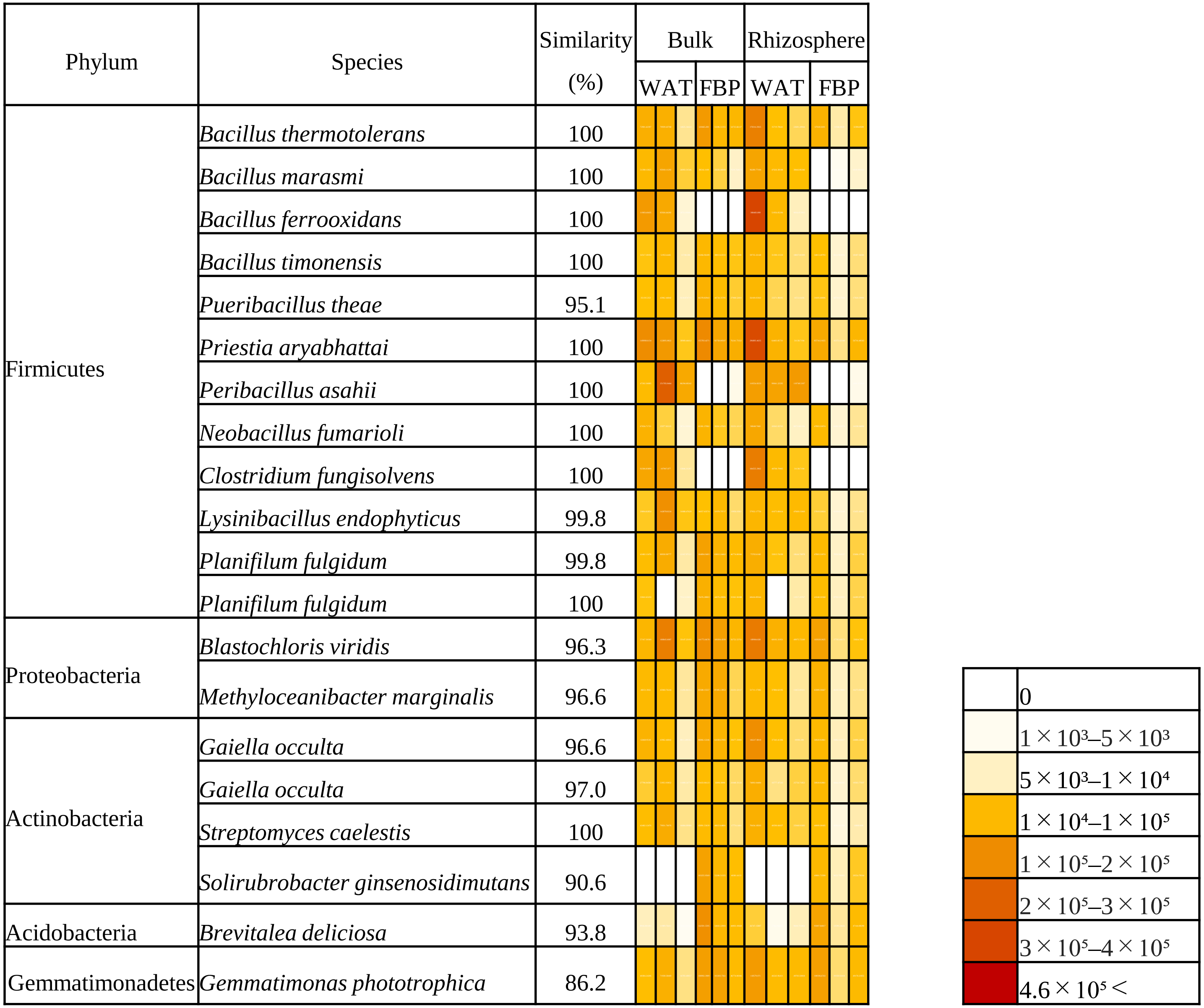


Fig. S13 Heatmap of predominant species copy number in bulk and rhizosphere soils applied with water (WAT) or 10,000-fold diluted FBP with after 28 days of komatsuna (*B. rapa)* cultivation.

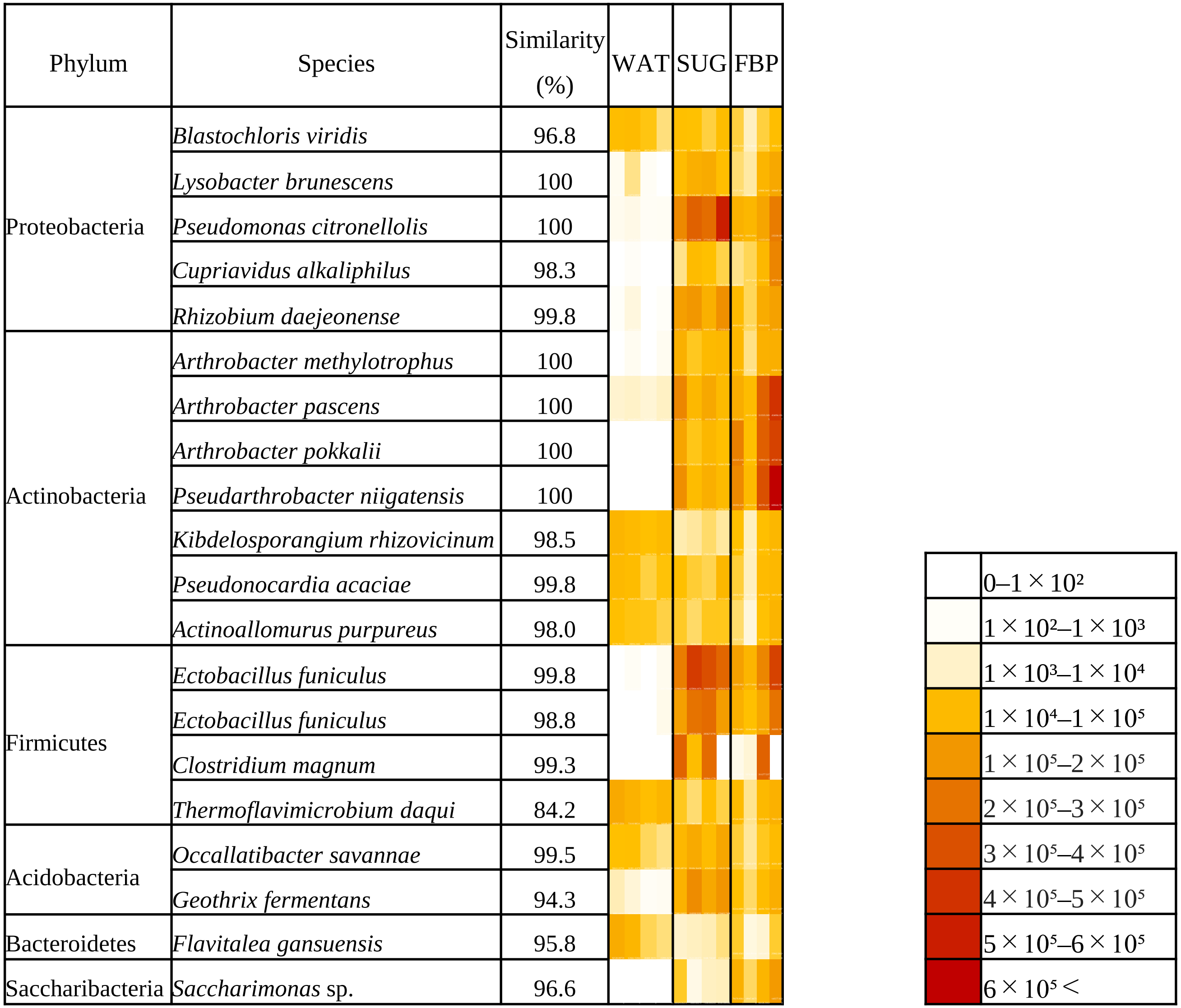


Fig. S14 Heatmap of predominant species copy number in soil applied with water (WAT), 200-fold diluted brown sugar (SUG) or 100-fold diluted FBP at 25° C for two weeks in a glass chamber.

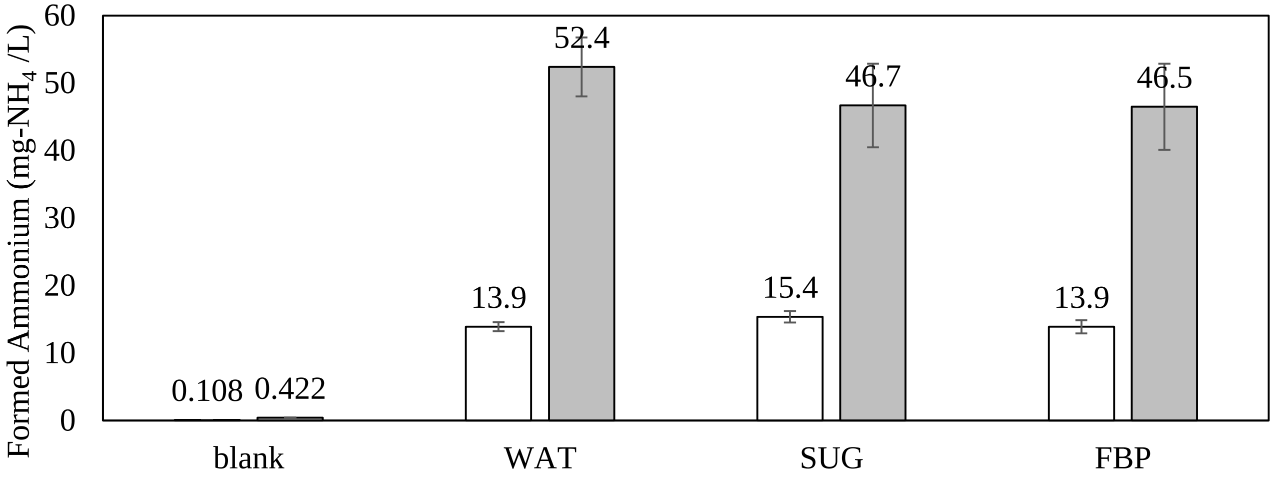


Fig. S15 Ammonium formation in soils of immersion test suspended with 0.1 g/L (white) and 0.5 g/L (black) peptone solution after 3 days incubation. Soil immersion test was performed applied with water (WAT), 200-fold diluted brown sugar (SUG) or 100-fold diluted FBP for two weeks in a glass chamber. Data are presented as the mean ± standard error bar (n=4). The statistical method used was Student t-test. No significant differences were obtained between water and FBP applications.

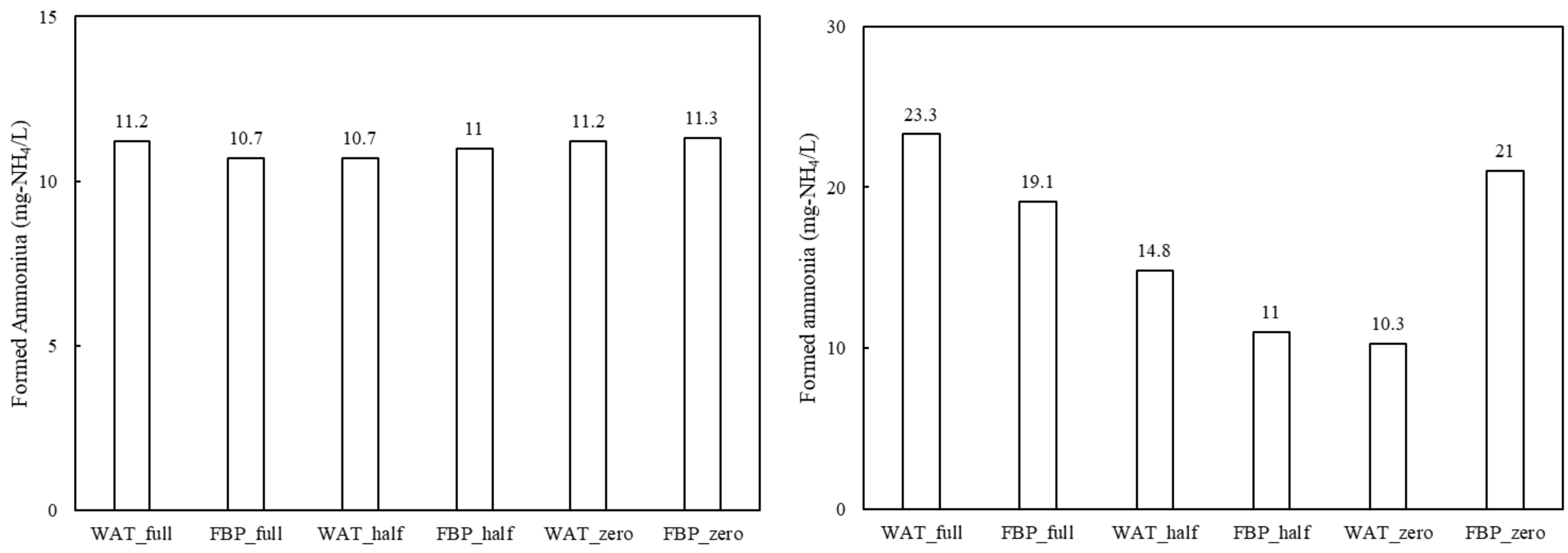


Fig. S16 Ammonium formation in soils of tomato cultivation suspended with 0.1 g/L (left) and 0.5 g/L (right) peptone solution after 3 days incubation. Tomato cultivation was performed applied with water (WAT) or 5,000-fold diluted FBP and chemical fertilizer dosages at 0% (zero), 50% (half), and 100% (full) for 112 days. Data are presented as the mean ± standard error bar (n=5). The statistical method used was Student t-test. No significant differences were obtained between water and FBP applications.

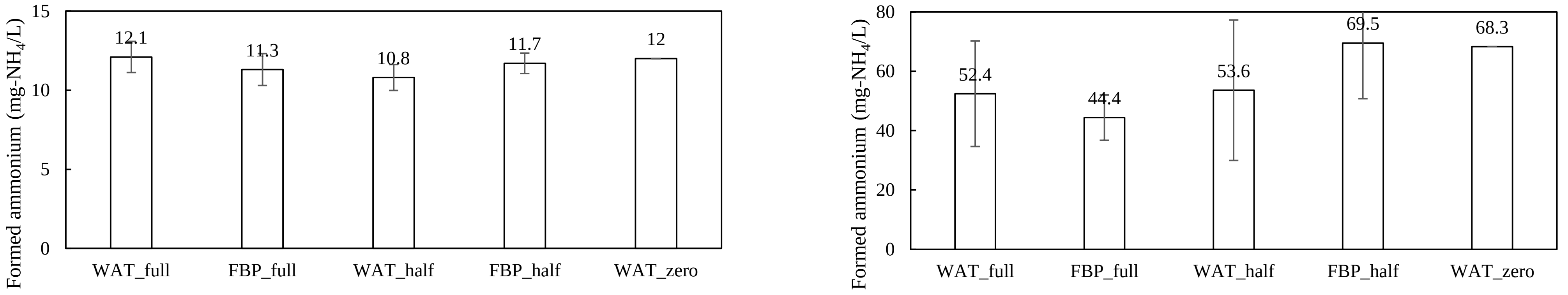


Fig. S17 Ammonium formation in soils of rice cultivation suspended with 0.1 g/L (left) and 0.5 g/L (right) peptone solution after 3 days incubation. Rice cultivation was performed applied with water (WAT) or 5,000-fold diluted FBP and chemical fertilizer dosages at 0% (zero), 50% (half), and 100% (full) for 129 days. Data are presented as the mean ± standard error bar (n=3). The statistical method used was Student t-test. No significant differences were obtained between water and FBP applications.

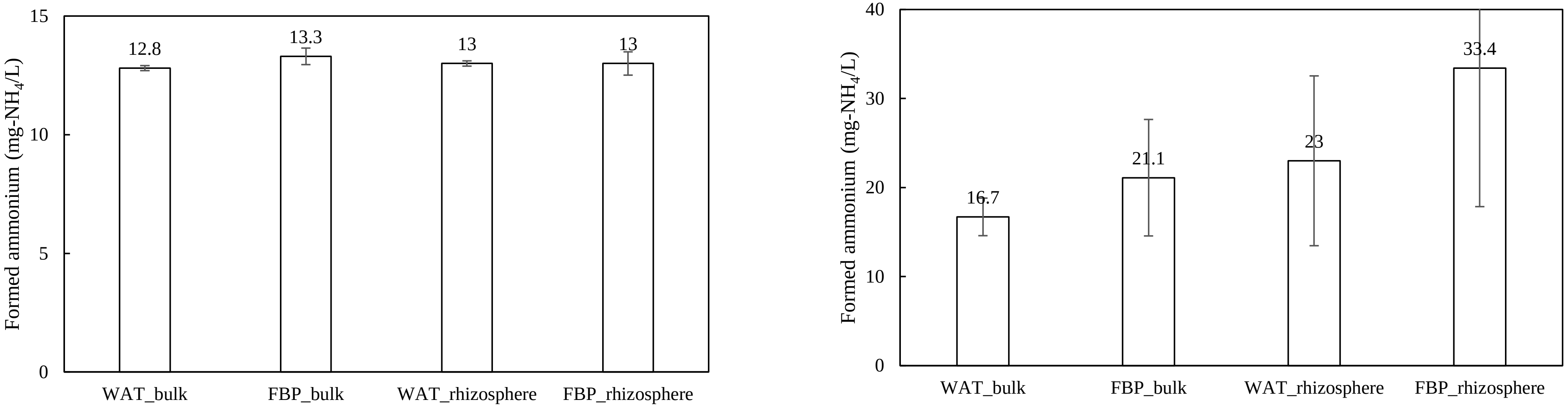


Fig. S18 Ammonium concentration in bulk and rhizosphere soils suspended with 0.1 g/L (left) and 0.5 g/L (right) peptone solution after 3 days incubation. Komatsuna (*B. rapa)* cultivation was performed applied with water (WAT) or 10,000-fold diluted FBP for 28 days. Data are presented as the mean ± standard error bar (n=3). The statistical method used was Student t-test. No significant differences were obtained between water and FBP applications.

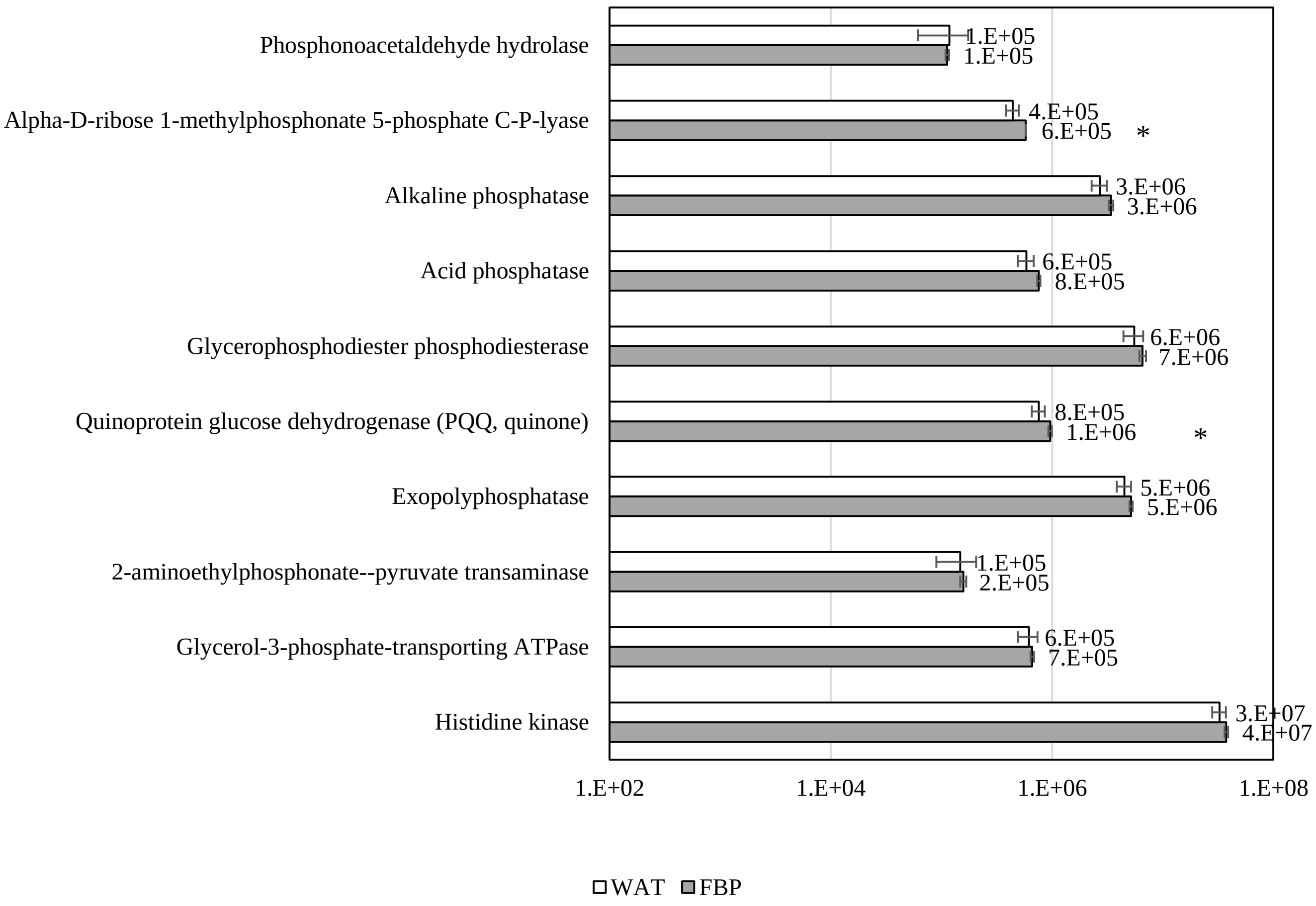


Fig. S19 The numbers of predicted genes related to phosphate solubilization in soil microbiota applied with water (WAT) or 5,000-fold diluted FBP after 129 days of rice cultivation (n=3). Asterisks indicate significant differences from WAT, as assessed by Student’s t-test: *$p < 0.05$.

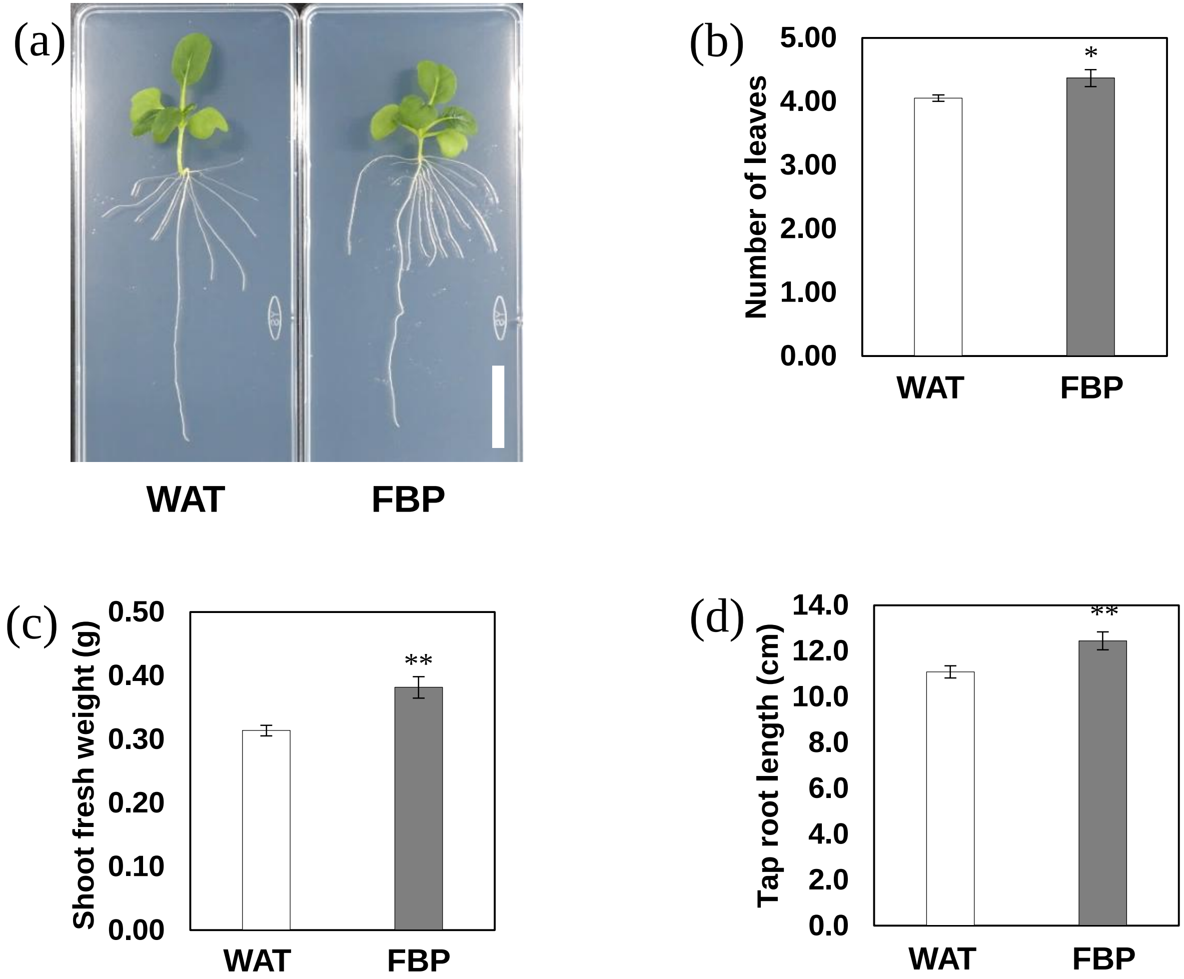


Figure S20. Effects of FBP on komatsuna (*Brassica rapa* var. *perviridis*) root elongation and growth development. Komatsuna was grown under 1/2 MS agar treated with water (WAT) or 2,500-fold diluted FBP (FBP). (a) Morphology of komatsuna after 21 day of cultivation. Bars=3 cm. (b)Number of leaves, (c) Shoot fresh weight, (d) Total root length after 21 day of cultivation. Data are presented as the mean ± standard error (n=19). Asterisks indicate significant differences from WAT, as assessed by Student's t-test: *$p < 0.05$, **$p < 0.01$.
